# Multiple layers of phospho-regulation coordinate metabolism and the cell cycle in budding yeast

**DOI:** 10.1101/815084

**Authors:** Lichao Zhang, Sebastian Winkler, Fabian Schlottmann, Oliver Kohlbacher, Josh E. Elias, Jan M. Skotheim, Jennifer C. Ewald

## Abstract

The coordination of metabolism and growth with cell division is crucial for proliferation. While it has long been known that cell metabolism regulates the cell division cycle, it is becoming increasingly clear that the cell division cycle also regulates metabolism. In budding yeast, we previously showed that over half of all measured metabolites change concentration through the cell cycle indicating that metabolic fluxes are extensively regulated during cell cycle progression. However, how this regulation is achieved still remains poorly understood. Since both the cell cycle and metabolism are regulated to a large extent by protein phosphorylation, we here decided to measure the phosphoproteome through the budding yeast cell cycle. Specifically, we chose a cell cycle synchronisation strategy that avoids stress and nutrient-related perturbations of metabolism, and we grew the yeast on ethanol minimal medium to force cells to utilize their full biosynthetic repertoire. Using a tandem-mass-tagging approach, we found over 200 sites on metabolic enzymes and transporters to be phospho-regulated. These sites were distributed among many pathways including carbohydrate catabolism, lipid metabolism and amino acid synthesis and therefore likely contribute to changing metabolic fluxes through the cell cycle. Among all one thousand sites whose phosphorylation increases through the cell cycle, the CDK consensus motif and an arginine-directed motif were highly enriched. This arginine-directed R-R-x-S motif is associated with protein-kinase A, which regulates metabolism and promotes growth. Finally, we also found over one thousand sites that are dephosphorylated through the G1/S transition. We speculate that the phosphatase Glc7/ PP1, known to regulate both the cell cycle and carbon metabolism, may play an important role because its regulatory subunits are phospho-regulated in our data. In summary, our results identify extensive cell cycle dependent phosphorylation and dephosphorylation of metabolic enzymes and suggest multiple mechanisms through which the cell division cycle regulates metabolic signalling pathways to temporally coordinate biosynthesis with distinct phases of the cell division cycle.

## Introduction

For cells to proliferate, they need to coordinate cell growth driven by metabolism with the cell division cycle, which ensures that DNA and other crucial cellular components are duplicated and divided between two daughter cells. In budding yeast, it was viewed that cell metabolism and growth proceed largely independently of the cell cycle. This assumption comes from the observation that mutants arrested in distinct phases of the cell cycle continued to grow and became extremely large and irregularly shaped (Hartwell et al., 1974; Johnston et al., 1977; Pringle and Hartwell, 1981). This showed clearly that a cell cycle arrest does not stop metabolism and mass accumulation, which led to the text book model that in budding yeast growth controls division, but not vice versa (Morgan, 2007).

While the hierarchy of metabolism driving the cell cycle was long the consensus, many studies over this past decade have challenged this view. It now seems that metabolism, growth and division are tightly and multi-directionally coordinated in all eukaryotes including yeast (Goranov and Amon, 2010; Ewald, 2018). Indeed, several core cell cycle regulators also target metabolic pathways and thereby control metabolism and growth: The most central cell cycle regulator, the cyclin-dependent kinase (CDK), has been found to directly target proteins in carbohydrate and energy metabolism in yeast (Ewald et al., 2016; Zhao et al., 2016), flies (Icreverzi et al., 2012) and mammals (Galbraith et al., 2017; Wang et al., 2017) (reviewed in (Solaki and Ewald, 2018)). Moreover, in addition to its role in mitosis, the polo kinase routes fluxes through the pentose-phosphate pathway by phosphorylating glucose-6-phosphate dehydrogenase in human cancer cell lines (Ma et al., 2017), and the cell cycle regulated ubiquitin ligase APC/C (anaphase promoting complex) regulates glucose metabolism in HeLa cells (Tudzarova et al., 2011). However, while specific examples of cell cycle regulators controlling metabolic pathways are accumulating, the global scope of metabolic regulation during the cell cycle is still largely unexplored.

The global regulation of metabolic processes during cell cycle progression is likely to be vast because 50% of the measured metabolites in budding yeast change concentration significantly in cells released synchronously into the cell cycle from a G1 arrest (Ewald et al., 2016). This suggests there are still many regulatory interactions coordinating metabolism and growth with cell cycle progression to be discovered. So far, we do not know which metabolic enzymes are targeted by which signalling pathways to control metabolic fluxes during the cell cycle.

To begin to address cell cycle-dependent regulation of metabolism, we performed a time-resolved proteome and phospho-proteome study through the cell cycle in synchronized yeast cultures. While there have been several phospho-proteomics reports on the budding and fission yeast cell cycle (Archambault et al., 2004; Holt et al., 2009; Carpy et al., 2014; Swaffer et al., 2016; Touati et al., 2018; Touati and Uhlmann, 2018), there are two important factors that make this study unique and complementary to previous work: First, we employed a synchronization strategy that releases cells from a G1 arrest without external perturbations of metabolism such as media switches, temperature shifts, addition of toxic chemical, or physical stress (Ewald et al., 2016; Rosebrock, 2017). Second, nearly all yeast cell cycle studies are performed using cells growing on complex or synthetic complete media, while we grow cells on ethanol minimal medium to force cells to activate a much larger repertoire of biosynthetic pathways. We found that more than two hundred phosphorylation sites on metabolic enzymes and transporters change in abundance during the cell cycle. Our data further suggests that metabolic signalling pathways including PKA, Snf1, and Glc7 are transiently regulated during cell cycle progression. Thus, we provide evidence for multiple layers of phospho-regulation that coordinate metabolism with cell cycle progression.

## Results

In this study, we wanted to identify mechanisms coordinating metabolism with cell cycle progression. Since both the cell cycle (Morgan, 2007; Enserink and Kolodner, 2010) and metabolic fluxes (Oliveira et al., 2012; Conrad et al., 2014; Chen and Nielsen, 2016) are known to be strongly regulated by phosphorylation, we decided to perform a phospho-proteomics and total proteomics time course of cells progressing through the cell cycle. Specifically, we arrested cells growing on ethanol minimal medium in G1 using our previously described hormone-inducible-cyclin strains (Ewald et al., 2016). These cells lack endogenous G1 cyclins (*cln1Δcln2cln3Δ*) and have an exogenous copy of *CLN1* that is expressed from an estradiol-inducible promoter (*LexApr-CLN1*) (Ottoz et al., 2014). Importantly, this strain can be released from a G1 arrest by adding 200 nM estradiol, which induces G1 cyclin expression without any other detectable cellular perturbations. Avoiding perturbations such as media changes, physical or temperature stress during the synchronous release is crucial when aiming to study metabolism, because many metabolic pathways are regulated in response to stress (Gasch and Werner-Washburne, 2002; Brauer et al., 2008). With this hormone-inducible strain, we performed two replicate experiments which showed very similar and highly synchronous budding profiles (Figure 1A-B). We note that we present data for the first two hours after the G1 release, which corresponds to most cells being in early mitosis and is before cells lose synchrony (Ewald et al., 2016).

**Figure 1:**
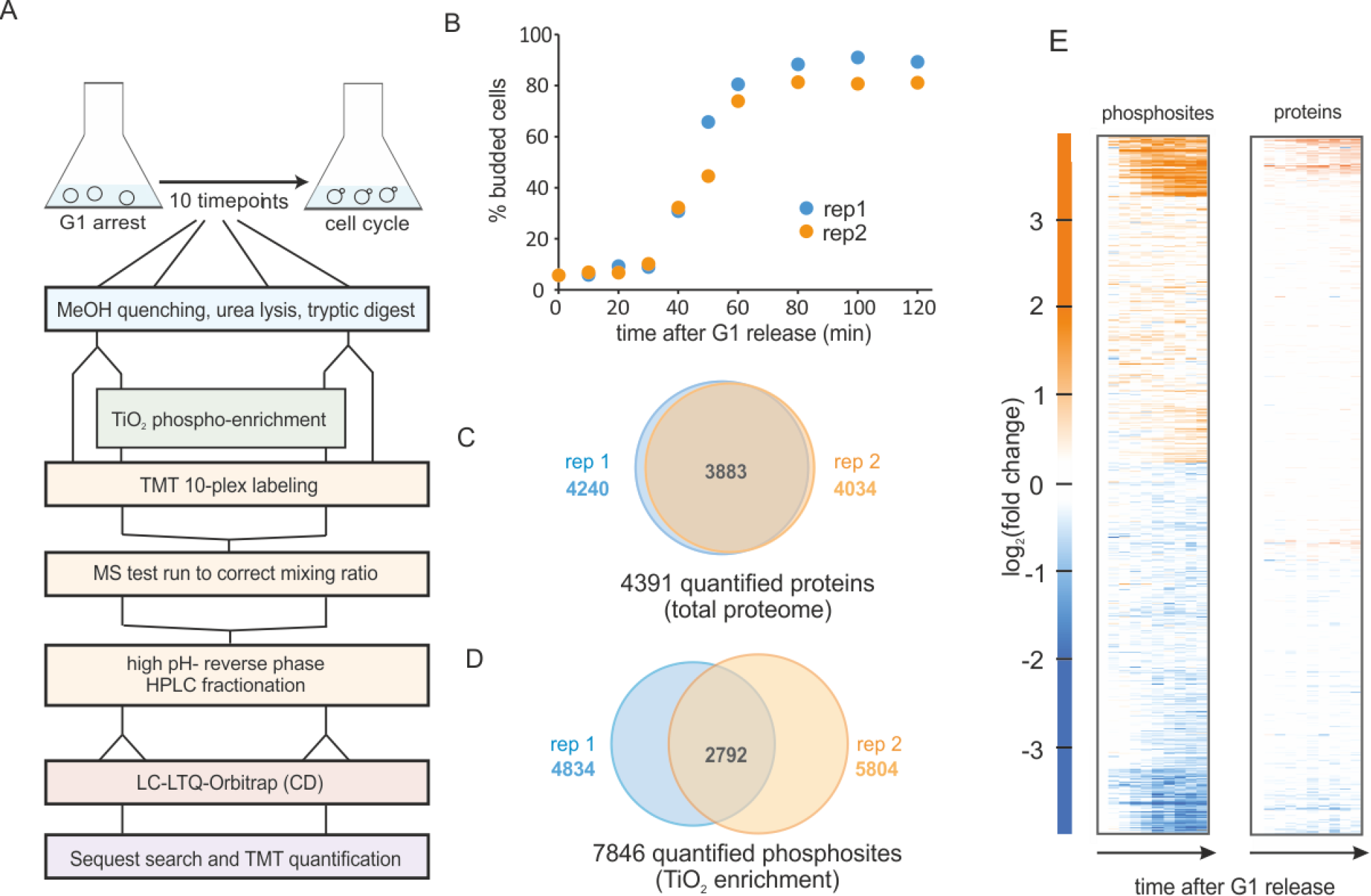
Phospho-proteomics time course of yeast cells released synchronously from G1 on ethanol minimal medium. A. Experimental workflow for sampling, phospho-enrichment, TMT labelling and mass spectrometry analysis B. Budding index of two replicate cultures released from a G1 arrest. C. Total protein and D. phosphorylated sites quantified in the two replicate experiments. E. Heatmap of the averaged replicates (log_2_ fold changes relative to t=0 min) for phosphorylated sites and quantified proteins.

From our two cell cycle synchronized cultures, we sampled ten time points from each replicate. Cells were lysed, proteins were digested with trypsin and lysC, and phosphopeptides were enriched with TiO_2_ and labelled with the TMT-10 plex (Figure 1A and methods).

In our total proteome cell cycle time course, we quantified over 4,000 proteins, with more than 90% overlap between the replicates (Figure 1C, Supplementary Table 1). Using an MS3 approach (25) and stringent quality criteria (see methods) we quantified a total of 9,267 unique phosphopeptides across all time points. This resulted in almost 8,000 quantified phosphorylation sites with approximately half of these quantified in both replicates (Figure 1D, Supplementary Table 2). As reported in previous studies (14, 26), the overall changes in the proteome through the cell cycle are small. In contrast, approximately one third of all phospho-sites change in abundance during the cell cycle suggesting cell cycle-dependent phosphorylation of these sites (Figure 1E).

Next, we sought to identify which phosphorylation sites where regulated during the cell cycle and test the quality and reproducibility of our phosphoproteome data. We first ranked the time profiles of all phosphorylation sites based on a heuristic p-value of change across the cell cycle (see methods). We then removed sites from further analysis that strongly correlated with total protein abundance, since these are unlikely to be regulated mainly by phosphorylation. We used the top third of the sites based on our ranking for further analysis (Supplementary Figure 1). To test the quality and reproducibility of our data, we correlated all ten time points of replicate 1 with all ten time points of replicate 2. Samples from corresponding times after release correlated well with p-values (Pearson correlation) of 10^−15^ or less for each of the ten time points (Figure 2A). As expected, neighbouring time points show a higher degree of correlation than more distant data points. Moreover, a principle component analysis (PCA) separated the samples according to the time they were taken along the first component, and replicate samples were positioned near each other in the first two PCA components (Figure 2B), an indication of the accuracy of the acquired data. To test if our data captures known cell cycle regulation, we used the DeRegNet software (Winkler et al in preparation, see methods), which identifies regulated subnetworks from large interaction networks. Here, we used the KEGG interaction network and searched for regulated subnetworks in our top-ranking phospho-sites (see methods). This approach recapitulated many aspects of the G1/S regulation (Figure 2C), indicating that our data is in good agreement with known cell cycle regulation.

**Figure 2:**
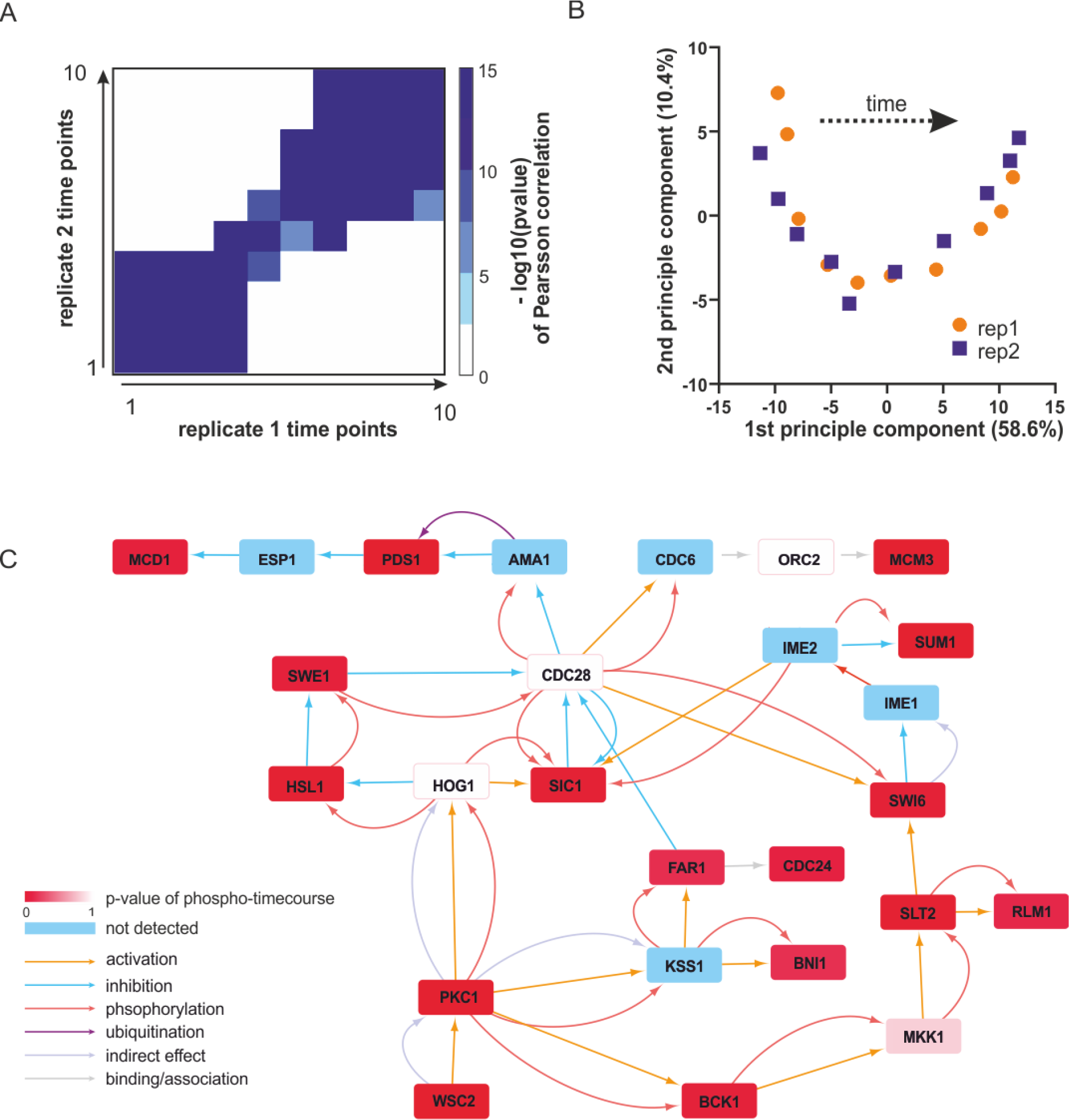
Data overview and quality controls. A. All time points of replicate 1 were correlated with all time points from replicate 2 (based on top 3^rd^ ranking phosphosites, see methods). Shown is a heatmap of the – log_10_(p-value) of a Pearsson correlation for all time points of one replicate with those of the other replicate. Principle component analysis performed with the top 3^rd^ of the identified phosphosites. Plotted are the ten time points of each replicate projected onto the first two principle components. C. Regulated sub-network identified from the top-ranking phosphoproteome data by the DeRegNet software (see methods) based on the KEGG interaction network. The type of interaction annotated in KEGG is indicated by the colour of the arrow.

Having established the quality of our phosphoproteomics time course, we next investigated which metabolic enzymes were dynamically phosphorylated and possibly regulated. To analyse the trends in the data set and how they relate to metabolism, we clustered the top-ranking sites using k-means clustering into five distinct clusters (Figure 3A-B; four, six and eight clusters give qualitatively similar results as shown in Supplementary Figure 2). For each cluster, we analysed which of the phosphorylation sites were annotated to proteins listed in the yeast metabolome database (Ramirez-Gaona et al., 2017) (Figure 3A). Proteins related to metabolism were found in every cluster, and, in total 243 sites on 134 metabolic proteins were changing (Figure 3B). Interestingly, more sites on these metabolic proteins were dephosphorylated than phosphorylated (Figure 3C). To determine which metabolic pathways were most likely affected by phospho-regulation, we sorted the 81 most dynamic sites on metabolic proteins from clusters 1, 2, and 5 into KEGG categories. All major metabolic pathways were represented and there was no particular category enriched relative to the whole dataset. In line with our previous metabolomics data showing that over half of ~500 measured metabolites change throughout the cell cycle (Ewald et al., 2016), these phosphoproteomics data suggest that global adaptations across metabolism are occurring during the cell cycle and are at least in part regulated by phosphorylation.

**Figure 3:**
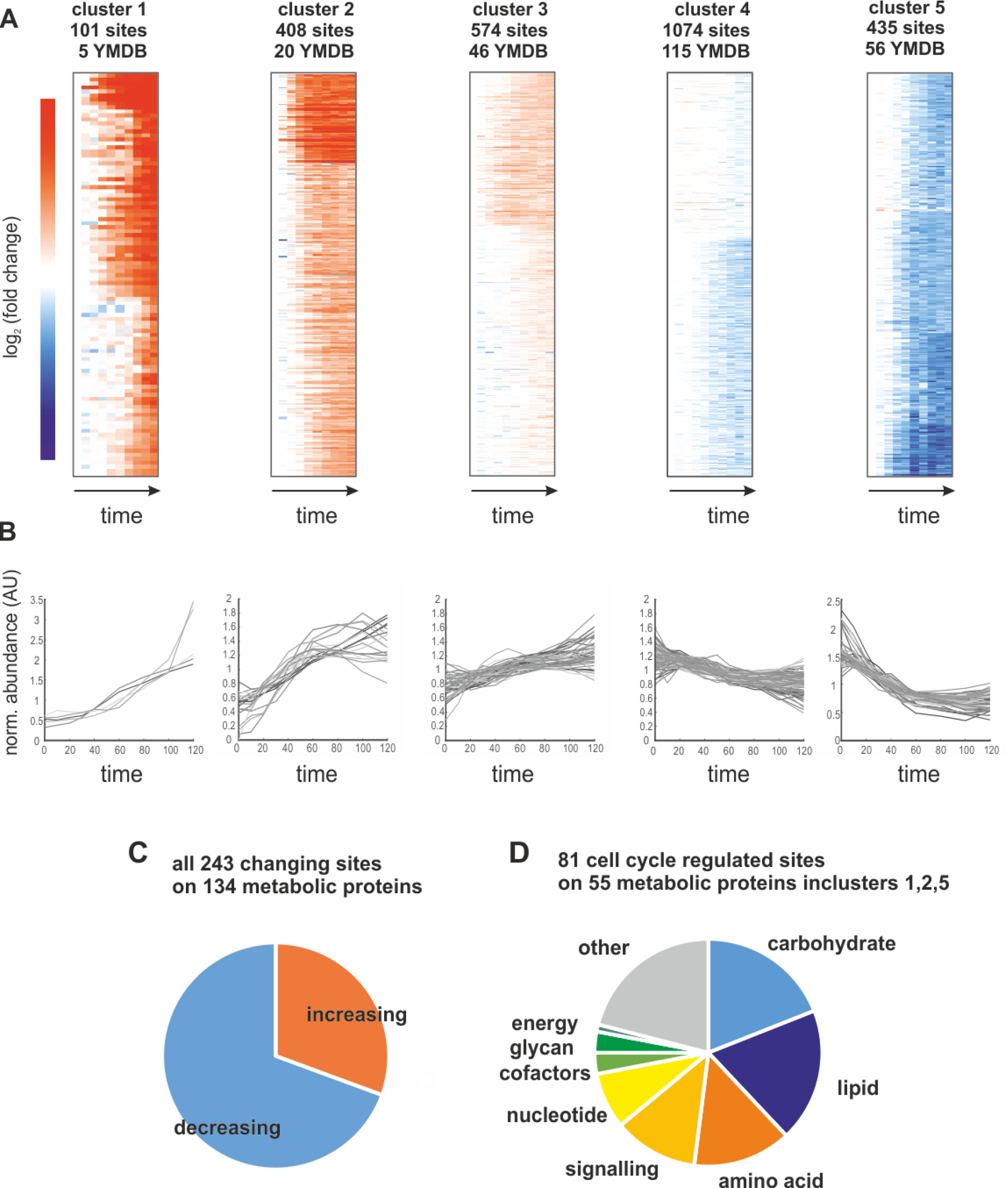
A. Heatmaps of the five identified clusters based on log_2_ (fold change) relative to t= 0 minutes. We report the number of sites contributing to the cluster and how many of those map to proteins in the yeast metabolome database (YMDB). B. Time course of phosphosite abundance for all sites on a YMDB protein in the corresponding cluster. C. Pie chart reporting the fraction of phosphosites on YMDB metabolic proteins whose abundance is increasing or decreasing through the cell cycle. D. Pie chart reporting the pathway assignments of most changing the phosphorylation sites whose abundance changes the most through the cell cycle.

We next wanted to determine which of the measured changes in enzyme phosphorylation may directly contribute to changes in metabolic activity. As a rough approximation of metabolic activity we use the product-to-substrate ratios from our previous metabolomics data set (Ewald et al., 2016). A change in the product-to-substrate ratio indicates a change in the kinetics of the reaction. For 174 sites on 82 proteins in our data set we had at least one substrate and one product (not including cofactors) for the reaction catalysed by the phosphorylated enzyme. For each of these reactions we correlated the phospho-site abundance with the product-to-substrate ratio (Supplementary Table 3). We found 19 sites on 15 enzymes with an R^2^ of the correlation greater than 0.5 (Supplementary Figure 3). One example is an enzyme well known to be upregulated during the cell cycle: the ribonucleotide-reductase complex, which catalyzes the conversion of NTPs to dNTPs (Lowdon and Vitols, 1973). The CDK consensus site S816 on Rnr1 correlates well with the ratio of dCTP to CTP (We note that cytosine nucleotides were chosen as example since they have unique masses in our metabolome data set and they do not participate in as many other reactions as adenylate or guanylate nucleotides) (Figure 4A-C). It therefore seems likely that Rnr1 S816 contributes to activating enzyme activity. Additionally, Rnr1 is also transcriptionally upregulated, but the increase in phosphorylation on S816 greatly exceeds the increase in total protein (Supplementary Figure 4). A second example is glutamine-fructose-6-phosphate amidotransferase (Gfa1), which catalyses the first step in the chitin pathway necessary for cell wall synthesis. The site S332 on this Gfa1 is dephosphorylated during the cell cycle which anti-correlates with the product to substrate ratio (Figure 4 D-F). We therefore suggest that this is an inhibitory phosphorylation which is being released during the cell cycle to increase chitin synthesis for surface expansion and cytokinesis. Whether this dephosphorylation is directly regulated by the cell cycle machinery or whether it is a secondary effect downstream of other metabolic changes (such as trehalose and glycogen utilization (Ewald et al., 2016; Zhao et al., 2016)) remains to be investigated. The resulting slopes and R^2^ of all correlations that could be determined based on the two datasets are reported in Supplementary Table 3.

**Figure 4:**
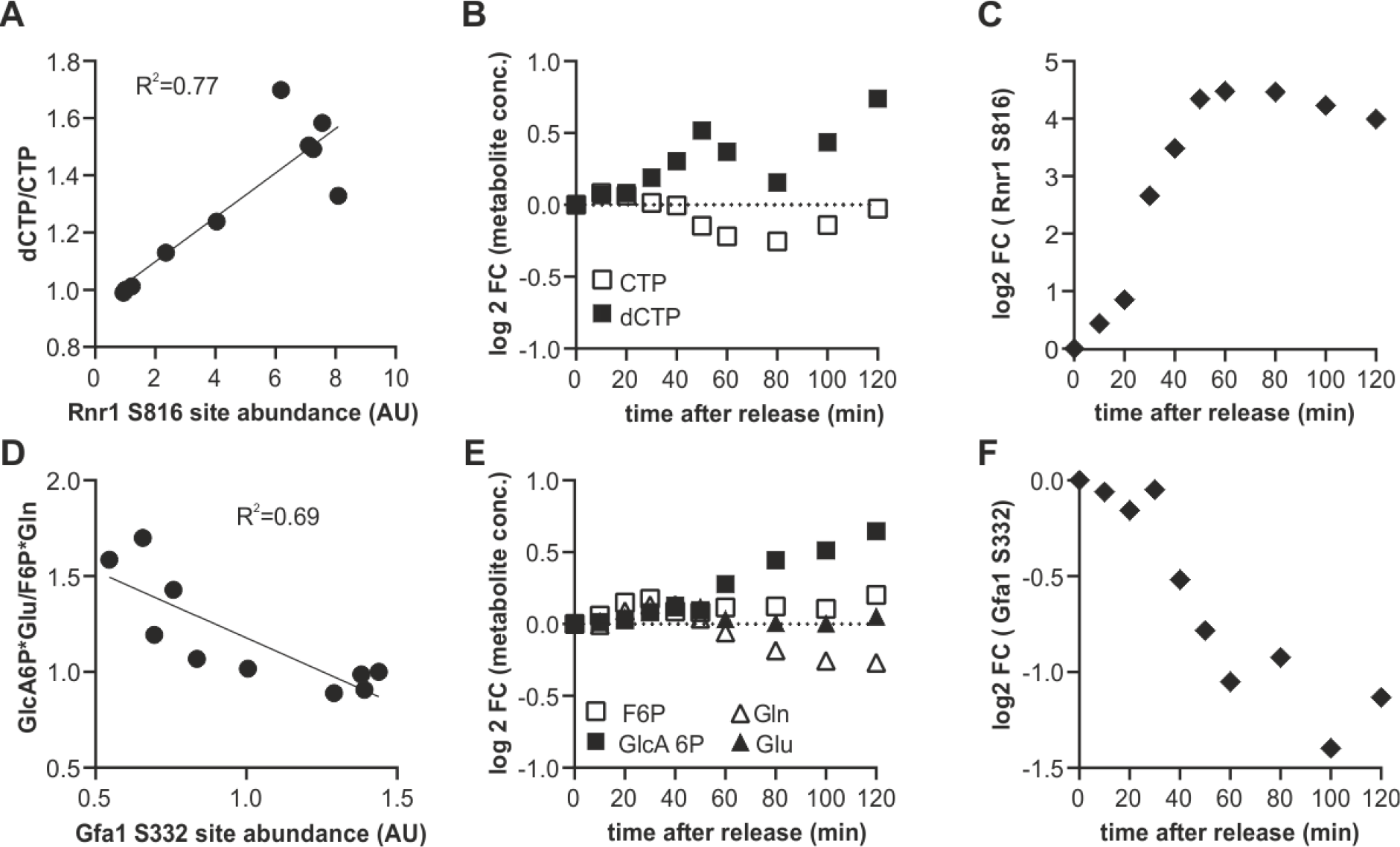
Identification of putative potential flux controlling phosphorylation sites based on the product to substrate ratio from published metabolomics data (6) A.-C. Example of a putative activating phosphorylation site showing the correlation of the serine 816 on ribonucleoreductase 1 (Rnr1) with the dCTP to CTP ratio, and the corresponding cell cycle time courses. D-F. Example of a putative inhibiting phosphorylation site showing anti-correlation of serine 332 of Glutamine-fructose-6-phosphate amidotransferase (Gfa1) with the ratio of its products and substrates.

To investigate which kinases contribute most to increasing phosphorylation in metabolic and all other proteins, we performed an unbiased motif analysis using the motif-x algorithm (Schwartz and Gygi, 2005) implemented on the Meme-suite (Bailey et al., 2009; Cheng et al., 2019). Not surprisingly, the two clusters corresponding to phosphorylation sites increasing early and late through the cell cycle were highly enriched for CDK consensus sites (S/T-P-X-K/R) and minimal CDK sites (S/T-P) sites (Figure 5A-B). However, the most enriched motif in the gradually increasing cluster 3 was RRxS/T and not proline-directed. This motif is the consensus sequence associated with the protein kinase A (PKA) and some other kinases (Ptacek et al., 2005; Mok et al., 2010). In clusters 1-3, which contained all sites increasingly phosphorylated through the cell cycle, almost half were proline directed and 15% were arginine directed (putative PKA targets) (Figure 5C). When we were only examining phosphorylation sites on metabolic proteins, we obtained a similar distribution (Figure 5D).

**Figure 5:**
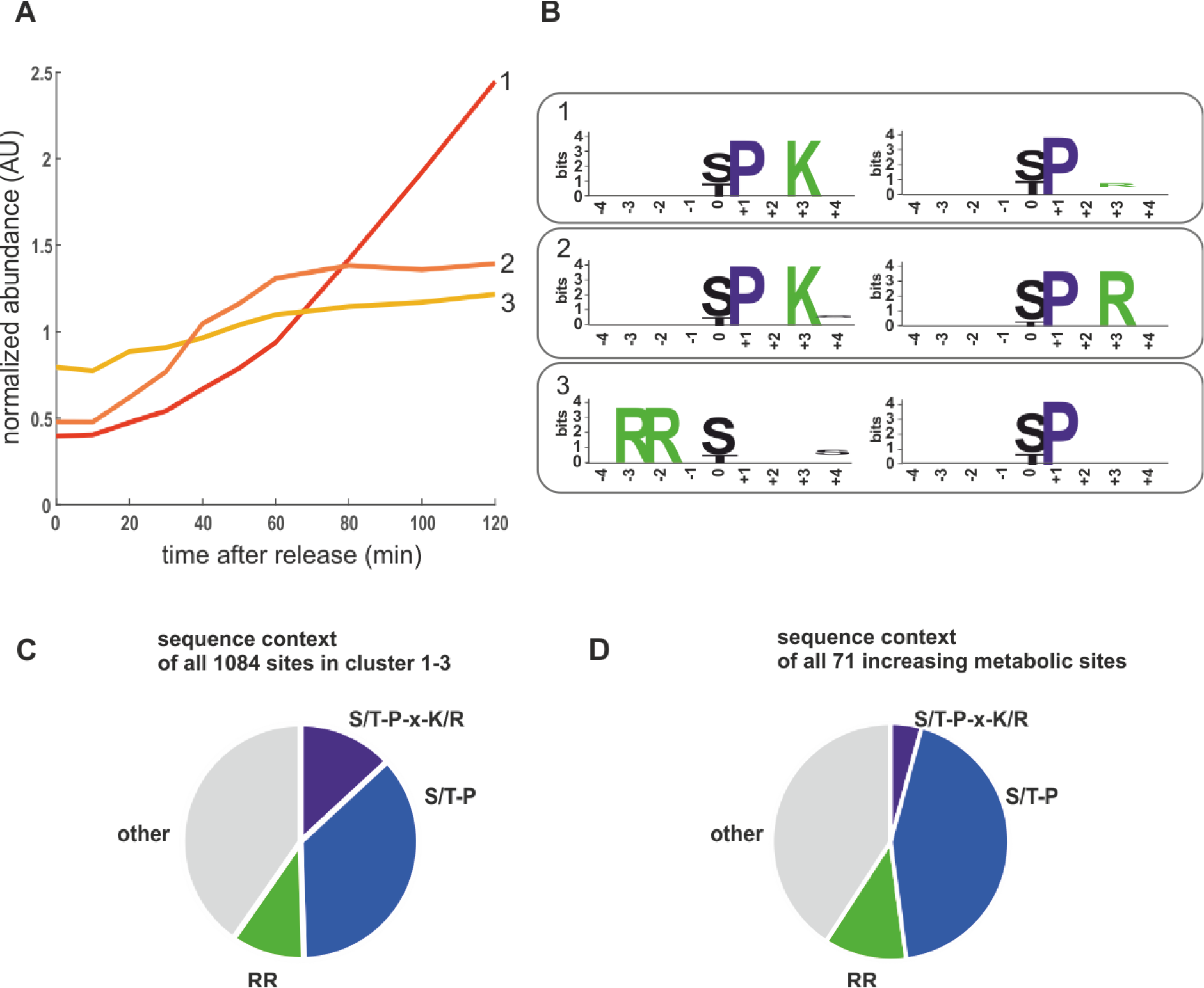
Motifs enriched in increasing phosphorylation sites. A. Cluster averages of three increasing clusters identified by kmeans clustering. B. Enriched motifs identified in the increasing clusters using the motif-X algorithm. The two most enriched motifs for each cluster are shown (>10-fold enriched, p< 10^−6^). Pie chart depicting the sequence context of all sites in the cell cycle increasing clusters 1-3. D. Pie chart depicting the sequence context of the cell cycle increasing phosphosites on metabolic proteins. RR denotes motifs potentially recognized by PKA including RRxS, RRxxS, and RxRxS. S/T-P-x-K/R is the optimal CDK consensus site.

That we identified consensus PKA phosphorylation sites as being dynamic through the cell cycle is interesting because PKA kinase is a sensor of nutrients (mainly glucose) and environmental stresses. PKA promotes cell growth and glucose repression and inhibits several stress responses (Broach, 2012; Conrad et al., 2014). Since we did not change the nutrient or stress conditions of our yeast cultures, we wanted to further investigate how putative PKA target sites could be increasingly phosphorylated during cell cycle progression. We noticed that several regulators upstream of PKA seemed to be phospho-regulated during cell cycle progression, with several phosphorylation sites either increasingly or decreasingly phosphorylated through the cell cycle (Figure 6). Many of the increasingly phosphorylated sites were proline directed (Figure 6B, D, E) and were similar to CDK consensus sites. This suggests that the Ras-branch of the PKA pathway could be activated by the cell cycle machinery to control downstream processes in metabolism and growth.

**Figure 6:**
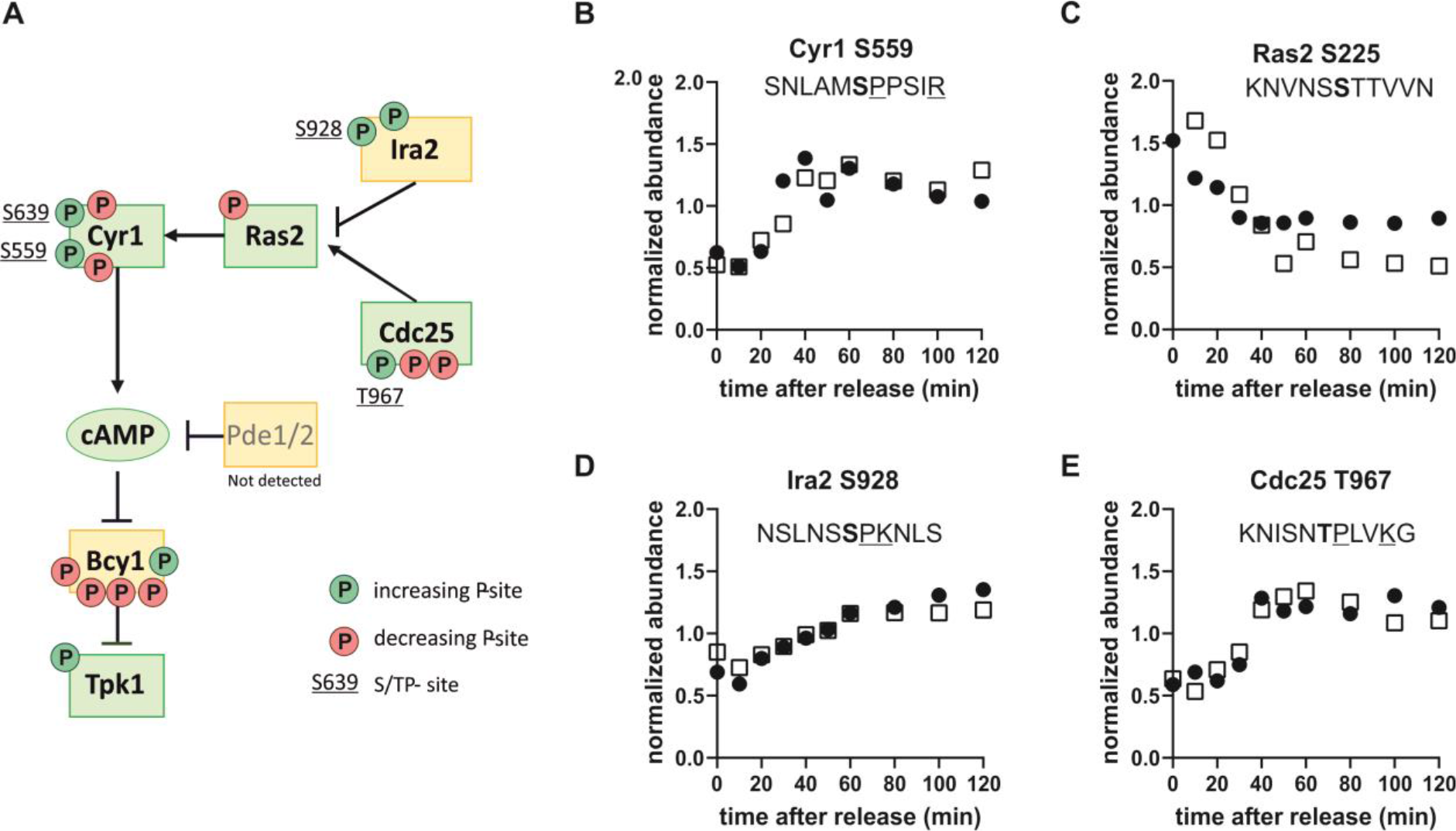
The protein kinase A pathway is phospho-regulated through the cell cycle. A. Map of the Ras-branch of the PKA pathway. Circles indicate sites whose phosphorylation increases (green, clusters 1-3) or decreases (red, clusters 4-5) through the cell cycle. Only sites found in both replicates are reported. S/TP sites, possibly phosphorylated by cyclin-dependent kinases, are denoted by their residue numbers adjacent to the phosphorylation site. B-E. Examples of dynamic phosphorylation of sites on different upstream regulators of PKA through the cell cycle. Residues associated with consensus cyclin-dependent kinase sites are underlined and the phosphorylated residue is shown in bold.

In addition to examining the sites increasingly phosphorylated through the cell cycle, we also wanted to investigate the sites being dephosphorylated through the cell cycle because they could be equally important. Dephosphorylation during the cell cycle is mainly discussed in the context of phosphatases counteracting CDK phosphorylation when cells go through mitosis (Mochida and Hunt, 2012; Rogers et al., 2016; Kataria et al., 2018) and in early G1 (Godfrey et al., 2017). In our experiment, we noticed that there are at least as many dephosphorylation events as phosphorylation events during the G1/S transition and S-phase, which are cell cycle transitions typically associated with increasing kinase activity. For metabolic proteins, twice as many sites were dephosphorylated through G1 to S as phosphorylated.

The prevalence of dephosphorylation through the cell cycle led us to wonder which phosphatases could be contributing, especially with regard to metabolism. In this context, we noticed that one of the top-ranking phosphorylation sites in our list was on Reg1, a regulatory subunit of the phosphatase Glc7 of the well-conserved PP1 family (Verbinnen et al., 2017). Glc7 has many targets and important functions in the cell cycle and in carbon metabolism (Cannon, 2010). Glc7 obtains its specific activity through interactions with regulatory subunits like Reg1 (Figure 7B) and has little specificity on its own. It does not seem to be regulated in abundance or in its phosphorylation state during the cell cycle (Supplementary Tables 1 and 2). Motivated by the identification of Reg1 as a dynamically phosphorylated protein, we searched our list of high-ranking phosphorylation sites for other Glc7 subunits. We found regulatory subunits that are known to regulate cell cycle functions including Bni4, which regulates bud neck and septum assembly, and Gip3, which regulates chromosome segregation (Figure 7A). Additionally, several of the subunits involved in regulating metabolism including Reg1 (glucose repression) and Gac1 (glycogen metabolism) were dynamically phosphorylated (Figure 7C). Although we did not find any annotated functions to these specific sites, it is tempting to speculate that these phosphorylation sites impact either binding of its targets or binding of the regulatory subunit to the catalytic subunit.

**Figure 7:**
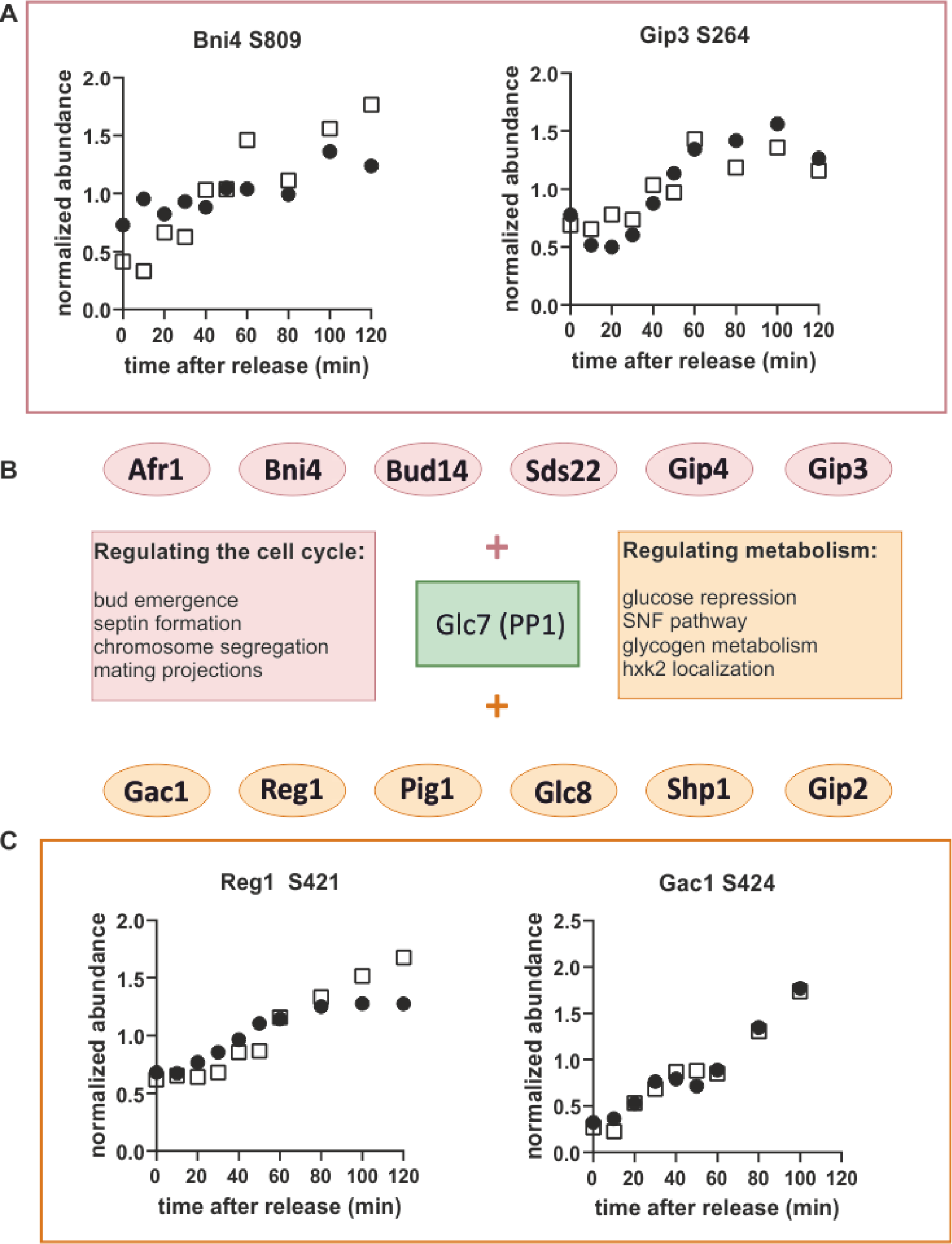
Glc7 (PP1) may regulate metabolism through the cell cycle. A. Cell cycle-dependent phosphorylation of the Glc7 subunits, Bni4 and Gip3, which are known to contribute to cell cycle regulation. B. Schematic showing regulatory subunits of the phosphatase Glc7 and their annotated functions. C. Cell cycle time courses of phosphorylation of the Glc7 subunits Reg1 and Gac1, which are known to contribute to metabolic regulation. Time courses from both replicates are shown.

To further investigate the idea that the cell division cycle drives changes in Glc7 phosphatase activity, we searched for known Glc7-Reg1 targets among our list of dephosphorylated sites. One of the most prominent targets of Glc7-Reg1 is the kinase Snf1 (homolog of mammalian AMPK (Hardie, 2011)). Snf1 is activated in the absence of glucose by phosphorylation on site T210 (Conrad et al., 2014). This activating phosphorylation is counter-acted by dephosphorylation by Reg1-Glc7 (Tu and Carlson, 1995). Consistent with our model, we find that Snf1 T210 is decreasing in abundance during the G1/S transition and seems to recover later in the cycle (Figure 8A). This was surprising given that Snf1 normally responds to changes in external glucose, which was constantly absent throughout our experiment. In response to glucose limitation, Snf1 regulates several aspects of carbon metabolism including the deactivation of the transcription factor Mig1. Mig1 is phosphorylated by Snf1 on at least four sites in its nuclear localization sequence and at least some of these sites are also reported to be dephosphorylated by Reg1-Glc7 (Smith et al., 1999). We therefore wondered whether Mig1 was also phospho-regulated during the cell cycle. We found one site S302, which closely follows the pattern of Snf1 dephosphorylation (Supplementary Figure 5A). While this site has not been specifically reported to be either a Snf1 or Reg1 target, it lies right between two Snf1 sites within the regulatory domain of Mig1 (Supplementary Figure 5C). Another site, T371, also lies within the Mig1 regulatory domain and is increasingly phosphorylated through the cell cycle (Supplementary Figure 5B). Interestingly, this site contains a proline in +1, which may point to phosphorylation by CDK1 as suggested by earlier studies (Holt et al., 2009; Zhao et al., 2016). A GFP-tagged Mig1 did not change localisation during the cell cycle under our growth conditions, suggesting these phosphorylation sites regulate Mig1 in a localisation-independent way.

**Figure 8:**
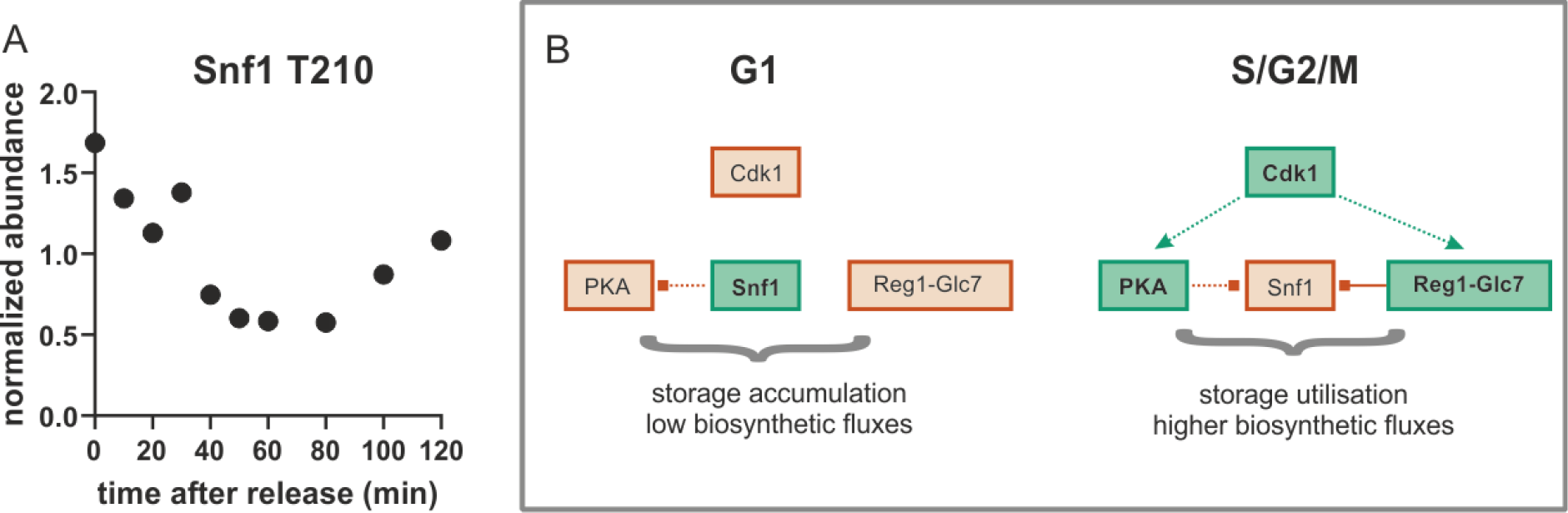
A. The well conserved activating site T210 on Snf1 is dephosphorylated during the G1-S-transition (average of both replicates) B. Model for global metabolic regulation during cell cycle progression on ethanol minimal medium. Red: low activity; green: higher activity; dotted lines: indirect or putative regulatory interactions; solid line: direct regulatory interaction

## Discussion

The aim of this study was to identify mechanisms coordinating metabolism and growth with the cell division cycle in budding yeast. Since both metabolism (Conrad et al., 2014; Chen and Nielsen, 2016) and the cell cycle (Morgan, 2008; Enserink and Kolodner, 2010) are extensively phospho-regulated, we performed a phosphoproteomics time-course of cells released from a G1 arrest. In contrast to previous phospho-proteomics studies, our main focus was to explore the phospho-regulation of metabolism through the cell cycle. We therefore took extreme care to employ a synchronisation strategy that would not lead to metabolic alterations through media changes or stress responses. To achieve this, our study was conducted with prototrophic strains growing on ethanol minimal medium, where cells grow slowly and need to activate their full biosynthetic potential. Our novel high quality dataset is therefore complementary to other phosphoproteomics data sets on the yeast cell cycle (Archambault et al., 2004; Holt et al., 2009; Touati et al., 2018; Touati and Uhlmann, 2018).

In summary, we found over 200 sites on metabolic enzymes that were either increasingly phosphorylated or dephosphorylated throughout the cell cycle. In agreement with our previous metabolomics study (Ewald et al., 2016), many different metabolic pathways were affected including carbohydrate, lipid, amino acid and nucleotide metabolism. While most of these sites still need to be functionally validated, the sheer number of phosphorylated or dephosphorylated sites suggests that phosphorylation contributes significantly to tailoring metabolic fluxes to the specific requirements of different cell cycle phases.

The identification of large-scale changes in phospho-isoforms through the cell division cycle raised the question as to which signalling pathways were responsible. We and others previously showed that the cyclin-dependent kinase directly regulates the activity of several metabolic enzymes such as the trehalase Nth1 (Ewald et al., 2016; Zhao et al., 2016) and the lipase Tgl4 (Kurat et al., 2009). This is unlikely to represent the full extent of metabolic regulation by CDK because previous work on rich media identified several other metabolic enzymes that were likely phosphorylated by CDK (Ubersax et al., 2003; Holt et al., 2009; Zhao et al., 2016). Using our minimal media conditions, we further expand the list of putative direct CDK targets in metabolism. However, the data also suggest that a direct regulation of enzymes by the cell cycle-dependent increase in proline directed CDK activity is not the main driver of adjusting metabolic fluxes, since many enzymes get dephosphorylated rather than phosphorylated, and only a minority of all phosphorylated sites are proline directed. We therefore suggest that a lot of the cell cycle-dependent phospho-regulation controlling metabolic fluxes is not directly through CDK activity, but entails additional pathways.

One such additional pathway could be the protein-kinase A signalling pathway. Our data suggests that the PKA pathway is cell cycle regulated and in turn contributes to cell cycle-dependent phosphorylation of downstream pathways. Two independent observations lead to this conclusion. First, the PKA consensus motif RRxS was found as highly enriched in one of the clusters of sites being increasingly phosphorylated through the cell cycle. Second, many of the upstream regulators in the Ras branch of the PKA pathway change in phosphorylation state during the early cell cycle. Many of these phosphorylation sites are proline directed, raising the possibility that CDK itself activates PKA signalling. If true, CDK regulation of PKA would provide a mechanistic explanation of the spikes in cyclic-AMP concentrations at the G1/S and G2/M transitions observed previously (Muller et al., 2003). Since PKA has been reported to regulate CDK activity at the G1/S transition (Tokiwa et al., 1994; Amigoni et al., 2015; Ewald, 2018), it is likely that the interplay between CDK and PKA is at the nexus coordinating metabolism, growth and division with nutrient supply.

While the putative PKA and CDK sites we identified are increasingly phosphorylated through the cell cycle, for many of the sites we identified the opposite is true. We were surprised at the large amount of dephosphorylation we observed as cells pass the G1/S transition. Many of these targets were metabolic enzymes. This large-scale dephosphorylation may be in part due to changing activity of the phosphatase Glc7/PP1 together with its subunits associated with metabolism such as Reg1 and Gac1. Reg1 also targets and inactivates another important metabolic signalling pathway such as the Snf1 Kinase, a member of the highly conserved AMPK family. Snf1 has a well characterized activating site T210 that is phosphorylated by upstream sugar sensing kinases and is dephosphorylated by Reg1. In both of our replicates, Snf1 T210 is dephosphorylated at the G1/S transition and re-phosphorylated as cells progress into mitosis consistent with the hypothesis that changing phosphatase activity may drive large-scale dephosphorylation through G1/S.

The dephosphorylation of Snf1 through G1/S may be important because when Snf1 is activated (like AMPK in mammals) it acts as a “brake pedal” slowing growth and energy consuming processes (Ghillebert et al., 2011; Coccetti et al., 2018). Thus, Snf1 inactivates many processes typically activated by PKA (Nicastro et al., 2015). During entry into the cell cycle at G1/S, phosphoregulation may shift the balance between PKA and Snf1 to enhance growth promoting pathways and rewire metabolism to turn storage compounds such as trehalose, glycogen or lipid droplets into macromolecules that support cell cycle progression (Figure 8B). This fine-tuned metabolic regulation likely does not matter much under the nutrient rich growth conditions (SCD, YPD) that most cell cycle studies are conducted in, but may be crucial in nutrient poor environments such as the ethanol minimal medium we used in this work.

Taken together, this and other work over the last decade (Kurat et al., 2009; Bryan et al., 2010; Goranov and Amon, 2010; Ewald et al., 2016; Zhao et al., 2016), shows that we need to revise the text book model that cell growth drives the cell cycle but not vice versa. Yeast physiology is likely determined by extensive cross talk between global regulators of metabolism, signalling pathways promoting growth, and the cell cycle control machinery (Ewald, 2018). More broadly, it seems safe to assume that all eukaryotes have extensive, multidirectional signalling mechanisms to coordinate metabolism, growth and the cell division cycle, given the many recent reports on the role of metabolism in proliferating tissues including cancer-, immune-, or stem cells (Vander Heiden and DeBerardinis, 2017; Corbet, 2018; Pearce and Pearce, 2018; Zhang et al., 2018; Dahan et al., 2019; Vaupel et al., 2019). We anticipate that over the coming decade this picture of interlinked metabolic and cell cycle control will be fleshed out as a broad array of post-translational modifications and allosteric interactions mediating cross-talk between metabolism and the cell division cycle are identified in model organisms and in humans.

## Author Contributions

J.C.E., J.M.S., and J.E.E. conceived and designed the study; J.C.E., L.Z., F.S. performed experiments; L.Z. and J.E.E. performed mass spectrometry and raw data analysis; S.W. and J.C.E. performed statistical analysis; O.K. advised on data analysis; J.C.E. and J.M.S. wrote the manuscript; all authors read and approved the manuscript.

## Acknowledgments

We thank Katja Kleemann for excellent technical support. We kindly acknowledge Boris Macek and his lab for helpful discussions.

## Funding

JCE gratefully acknowledges support by the Institutional Strategy of the University of Tübingen (Deutsche Forschungsgemeinschaft ZUK 63) We further acknowledge Deutsche Forschungsgemeinschaft and Open Access Publishing Fund of University of Tübingen for the support of publication costs. JMS was supported by the NIH through GM115479 and an HHMI-Simons Faculty Scholar Award.

## Conflict of Interest

The authors declare no conflict of interest.

## Data Accessibility

Processed data have been included as Supplementary Tables. The raw mass spectrometry proteomics data have been deposited to the ProteomeXchange Consortium via the PRIDE (Perez-Riverol et al., 2019) partner repository under the dataset identifier PXD015235.

## Materials and Methods

### Cell cultivation and synchronization

Cells were grown in 1 % ethanol minimal media (1.7 g yeast nitrogen base, 5 g/L ammonium phosphate, 10 ml ethanol, pH adjusted to 5 with potassium hydroxide) at 30 °C and 250 rpm orbital shaking. For cell cycle arrest strain JE 611c (Ewald et al., 2016) was grown on 10 nM estradiol to an OD of approximately 0.2. Cells were filtered, resuspended in estradiol-free medium, and grown for 15 hours. These G1 arrested cells were released by addition of 200 nM estradiol (dissolved at 1 mM in 100% ethanol). Cell cycle release was monitored by manual bud counting (>200 cells per sample) at 60x magnification.

### Sampling, protein extraction and digestion

20 ml of cell culture (OD ~0.6) were sampled into 1.5 volumes of 60% methanol and precooled to −40°C to quench metabolic activity. Cells were spun at 4000 g. The pellets were frozen in liquid nitrogen and then stored at −80°C until further use. Cells were lysed by bead beating in 8M urea, 150 mM NaCL, 5 mM DTT, 50 mM HEPES pH 8 supplemented with 1x Halt™ Protease and Phosphatase Inhibitor Cocktail (ThermoFisher Scientific). The lysate was centrifuged at 13,200 rpm for 15 min and the supernatant was transferred to fresh test tubes for a second round of centrifugation. Lysates from two parallel samples were combined to increase starting material. This was followed by an alkylation step using 14 mM iodoacetamide for 45 minutes at room temperature in the dark and the reaction was then quenched with DTT. In order to clean the proteins a methanol-chloroform precipitation was performed and the protein pellet was washed twice with acetone. The pellet was re-suspended with 8M urea in 50 mM HEPES (pH 8) and the total protein concentration was determined using the Pierce™ BCA Protein Assay Kit (Pierce, Rockford, IL). Approximately 4 mg of protein of each sample were diluted to 4 M urea using 50 mM HEPES (pH8) and digested with LysC (1:100) for 4 hours at room temperature. Samples were further diluted to 1 M urea using 50 mM HEPES (pH8) and trypsin (Promega, Madison, WI) was added at a ratio of 1:20 enzyme: substrate for 16 hours at 37 °C. The digestion was quenched with formic acid and the peptides desalted using a Sep-Pak C18 1 cc Vac 50 mg Cartridge (Waters, Milford, MA). 5% of each sample was used for total proteome analysis and the remaining peptide was used for phosphopeptide enrichment.

### Phosphopeptide enrichment

TiO_2_ powder was resuspended in 2M lactic acid/50% acetonitrile (binding solution) at a concentration of 25 mg/mL. Peptides were resuspended in 400 μl of binding solution and added to 640 μl of TiO_2_ slurry and incubated for one hour while shaking. The samples were then spun down at 10,000 rpm for 1 min and the supernatant was removed. The TiO_2_ pellet was washed with binding solution twice and then 0.1% trifluoroacetic acid/50% acetonitrile three times. Phosphopeptides were eluted off TiO_2_ using 50 mM KH_2_PO_4_ (pH 10 adjusted with ammonium hydroxide) twice, acidified with formic acid, and desalted using a Sep-Pak C18 column as above.

### TMT labelling and high-pH reversed-phase fractionation

The TMT labelling reagents were obtained from Pierce and the labelling was performed according to the manufactures suggested procedure and previously published protocol (Zhang and Elias, 2017). In brief, in brief, 100 μg samples were resuspended in 100 μl of 50mM Na-HEPES and then 30 μl of acetonitrile was added to each sample. A TMT-10plex kit was used and each TMT reagent (0.8 mg per vial) was reconstituted in 40 μl of acetonitrile. 10 μl of the reagent was added to the corresponding sample to incubate for 1 h. To reverse unwanted TMT labelling with tyrosine residues, the reaction was quenched with a final concentration of 0.3 % (v/v) hydroxylamine for 15 min at room temperature. Samples were acidified with formic acid to pH 2. In order to assess the labelling efficiency a ratio-check was performed by combining 5 μL of each sample, desalting by StageTip and then analysing with LC-MS. Based on the result from the ratio-check equal amounts of each individual labelled sample were then combined to deliver an overall equal amounts across all channels. The combined peptides were desalted using a Sep-Pak C18 column and then fractionated by high-pH reverse phase fractionation (Yang et al., 2012) using an 84 min gradient (buffer A: 10 mM ammonium formate, pH 10; buffer B: 10 mM ammonium formate, 90 % ACN, 10 % H_2_O, pH 10) on an Agilent 1200 HPLC (Agilent Technologies, Santa Clara, USA). In total 84 fractions were collected, concatenated, combined into a total of 12 fractions, and then dried down. All fractions were desalted using Sep-Pak C18 column, dried down and resuspended in 0.1% formic acid for LC-MS analysis.

### Mass Spectrometry Analysis

Peptides were separated on a 24 cm reversed phase column (100 μm inner diameter, packed in-house with ReproSil-Pur C18-AQ 3.0 m resin, Dr. Maisch GmbH) over 180 min using a two-step linear gradient with 4–25 % buffer B (0.2% (v/v) formic acid in acetonitrile) for 120 min followed by 25-45 % buffer B for 15 min at a 400 nL/min flowrate on an Dionex Ultimate 3000 LC-system (Thermo Scientific, San Jose, CA). Eluted peptides were analysed with a Fusion Lumos mass spectrometry system (Thermo Scientific, San Jose, CA). Full MS scans were performed in the Orbitrap in the mass range of 400-1500 m/z and the resolution was set to 120,000. The AGC setting was 4E5 and maximum injection time for FTMS1 was 50 ms. Data dependent mode was set to top speed with duty cycle of 3s. Precursor ions with charge states 2-7 were selected for fragmentation using collision induced dissociation (CID) with quadrupole isolation, isolation window of 0.7 m/z, normalized collision energy of 35% and activation Q of 0.25. MS2 fragments were analysed in the ion trap mass analyzer with turbo scan rate and maximum injection time of 50ms. Ions within a +/−10 ppm m/z window around ions selected for MS2 were excluded from further selection for fragmentation for 90 s. Following each MS2 CID, a MS3 higher-energy collisional dissociation (HCD) is performed with synchronous precursor selection enabled (the number of precursors set to 5) and collision energy of 65% (McAlister et al., 2014). HCD fragment ions were detected in the Orbitrap in the scan range of 120-500 m/z with resolution of 60,000, AGC setting of 10,000, and maximum ion time of 120 ms. The mass spectrometry proteomics data have been deposited to the ProteomeXchange Consortium via the PRIDE (Perez-Riverol et al., 2019) partner repository under the dataset identifier PXD015235.

### Data processing

#### Protein identification and quantification

Raw data were searched using SEQUEST in Proteome Discoverer 2.2 against a sequence database of yeast (strain W303, NCBI taxonomy ID 559292, downloaded on July 28, 2016). Trypsin was selected as the enzyme with at most two missed cleavage sites. Precursor mass tolerance was set to +/− 10 ppm and fragment mass tolerance was set to +/− 0.6 Da. At most three dynamic modifications were allowed per peptide. Carbamidomethylation of cysteine (+57.021 Da) and TMT-labelled N-terminus and lysine (+229.163) were set as static modifications. Oxidation of methionine (+15.995 Da) and acetylation of protein N-terminus (+42.011 Da) were set as variable modifications. For phosphopeptides analysis phosphorylation of Serine, Tyrosine and Threonine (+79.967) were also set as differential modifications. Percolator was applied to filter incorrect identifications down to an estimated false discovery rate of 1% for both peptides and proteins. The PtmRS node was used for phosphosite assignment. For quantification, a mass tolerance of +/−20 ppm window was applied to the integration of report ions using the ‘most confident’ centroid method and S/N values were reported as reporter abundances. For total proteome analysis, the threshold for average reporter S/N was set to 5, the threshold for co-isolation was set to 30%, and quantification results were rejected for missing channels. The data normalization mode was set to “total peptide amount” and scaling mode was set to “on channels average”.

#### Phosphorylation site quantification

For phosphosite analysis, PSMs were filtered to meet the following criteria: The phosphosite position confidence (ptmRS score) was set to > 75%; the threshold for average reporter S/N was set to 10; and the threshold for co-isolation was set to 30%. Only PSMs quantified in nine consecutive channels were included (so only the first or last time point were allowed to be zero). After filtering, the channels were normalized to the total intensity. PSMs were summed to unique peptides. Each phosphorylated site was then summed across all peptides containing that site. The quantification of each site was scaled by its mean before averaging the replicates.

### Statistical analysis

#### Heuristic p-value and ranking

To avoid any a priori assumptions of the shape of the time profiles, we ranked our time courses based on a heuristic p-value calculated in the following ways. For each phosphorylation site, we calculated a p-value from a t-test comparing the average of the first four to the last four time points. Also a regression over all timepoints as independent variables was performed to detect linear trends. Finally, we calculated the p-value of linear regressions in time windows of five time points moving across the time series to detect trends which do not span the whole time span. All values were corrected for multiple hypothesis testing with the Holm-Sidak correction. The minimum p-value obtained from these tests was then used to rank the phosphorylation sites.

To test whether this ranking separates changing from non-changing sites, we performed k-means clustering (see below) on sets of 1,000 sites from top to bottom rank, see Supplementary Figure 1. Based on the results from this clustering, we empirically decided to use the top third ranking phosphorylation sites for further analysis. For each site in each replicate, the correlation between the protein und phosphosite abundance was calculated. Phosphosites that correlated with Pearson’s R greater than 0.8 in either replicate were removed from downstream phosphorylation analysis. Above procedures were carried out with statsmodels (0.9.0) in Python 3.6.8.

#### K-means Clustering

k-means clustering was performed using the Matlab 2018b built-in algorithm with 1,000 iterations and 100 replicates. The number of clusters was empirically set to five (see Supplementary Figure 2 for results for 4, 6, and 8 clusters).

#### Principal Component Analysis

A principal component analysis was performed on the normalized abundance data using Perseus 1.6.1.3 (Tyanova et al., 2016).

#### Motif Enrichment

Motif enrichment was performed using the MoMo function (Cheng et al., 2019) on the MEME suite (http://meme-suite.org/, accessed in May 2019) (Bailey et al., 2009) with the following settings: motif-x algorithm; background peptides extracted from reference sequence GeneBank Saccharomyces cerevisiae uid 128; motif width 13; central residues with same modification mass combined; p-value threshold was set to 0.0001.

#### Subgraph Analysis

Deregulated subgraph were calculated with DeRegNet (https://github.com/sebwink/deregnet) (Winkler et al in prep, (Backes et al., 2012). DeRegNet takes a regulatory network (e.g. constructed from KEGG) and assigns a “deregulation” score to each node (protein) in the network. For every protein the minimum p-value across all associated sites was taken as a basis to calculate deregulation socres. As deregulation score we used binary scores defined as 1 for p-values < 0.1 and as 0 otherwise. DeRegNet then calculated a connected subnetwork within the Yeast KEGG network with maximal average deregulation score (sum of deregulation score of nodes in subgraph divided by number of nodes in the subgraph).

## References

Amigoni, L., Colombo, S., Belotti, F., Alberghina, L., and Martegani, E. (2015). The transcription factor Swi4 is target for PKA regulation of cell size at the G1 to S transition in Saccharomyces cerevisiae. Cell Cycle 14(15),2429–2438. doi: 10.1080/15384101.2015.1055997.

Archambault, V., Chang, E.J., Drapkin, B.J., Cross, F.R., Chait, B.T., and Rout, M.P. (2004). Targeted proteomic study of the cyclin-Cdk module. Mol Cell 14(6),699–711. doi: 10.1016/j.molcel.2004.05.025.

Backes, C., Rurainski, A., Klau, G.W., Muller, O., Stockel, D., Gerasch, A., et al. (2012). An integer linear programming approach for finding deregulated subgraphs in regulatory networks. Nucleic Acids Research 40(6). doi: ARTN e4310.1093/nar/gkr1227.

Bailey, T.L., Boden, M., Buske, F.A., Frith, M., Grant, C.E., Clementi, L., et al. (2009). MEME SUITE: tools for motif discovery and searching. Nucleic Acids Res 37(Web Server issue),W202–208. doi: 10.1093/nar/gkp335.

Brauer, M.J., Huttenhower, C., Airoldi, E.M., Rosenstein, R., Matese, J.C., Gresham, D., et al. (2008). Coordination of growth rate, cell cycle, stress response, and metabolic activity in yeast. Mol Biol Cell 19(1),352–367. doi: 10.1091/mbc.e07-08-0779.

Broach, J.R. (2012). Nutritional control of growth and development in yeast. Genetics 192(1),73–105. doi: 192/1/73 [pii]10.1534/genetics.111.135731.

Bryan, A.K., Goranov, A., Amon, A., and Manalis, S.R. (2010). Measurement of mass, density, and volume during the cell cycle of yeast. Proceedings of the National Academy of Sciences of the United States of America 107(3),999–1004. doi: 10.1073/pnas.0901851107.

Cannon, J.F. (2010). Function of Protein Phosphatase-1, Glc7, in Saccharomyces cerevisiae. Advances in Applied Microbiology, Vol 73 73,27–59. doi: 10.1016/S0065-2164(10)73002-1.

Carpy, A., Krug, K., Graf, S., Koch, A., Popic, S., Hauf, S., et al. (2014). Absolute Proteome and Phosphoproteome Dynamics during the Cell Cycle of Schizosaccharomyces pombe (Fission Yeast). Molecular & Cellular Proteomics 13(8),1925–1936. doi: 10.1074/mcp.M113.035824.

Chen, Y., and Nielsen, J. (2016). Flux control through protein phosphorylation in yeast. Fems Yeast Research 16(8). doi: ARTN fow09610.1093/femsyr/fow096.

Cheng, A., Grant, C.E., Noble, W.S., and Bailey, T.L. (2019). MoMo: discovery of statistically significant post-translational modification motifs. Bioinformatics 35(16),2774–2782. doi: 10.1093/bioinformatics/bty1058.

Coccetti, P., Nicastro, R., and Tripodi, F. (2018). Conventional and emerging roles of the energy sensor Snf1/AMPK in Saccharomyces cerevisiae. Microbial Cell 5(11),482–494. doi: 10.15698/mic2018.11.655.

Conrad, M., Schothorst, J., Kankipati, H.N., Van Zeebroeck, G., Rubio-Texeira, M., and Thevelein, J.M. (2014). Nutrient sensing and signaling in the yeast Saccharomyces cerevisiae. Fems Microbiology Reviews 38(2),254–299. doi: 10.1111/1574-6976.12065.

Corbet, C. (2018). Stem Cell Metabolism in Cancer and Healthy Tissues: Pyruvate in the Limelight. Frontiers in Pharmacology 8. doi: ARTN 95810.3389/fphar.2017.00958.

Dahan, P., Lu, V., Nguyen, R.M.T., Kennedy, S.A.L., and Teitell, M.A. (2019). Metabolism in pluripotency: Both driver and passenger? J Biol Chem 294(14),5420–5429. doi: 10.1074/jbc.TM117.000832.

Enserink, J.M., and Kolodner, R.D. (2010). An overview of Cdk1-controlled targets and processes. Cell Div 5,11. doi: 10.1186/1747-1028-5-11.

Ewald, J.C. (2018). How yeast coordinates metabolism, growth and division. Curr Opin Microbiol 45,1–7. doi: 10.1016/j.mib.2017.12.012.

Ewald, J.C., Kuehne, A., Zamboni, N., and Skotheim, J.M. (2016). The Yeast Cyclin-Dependent Kinase Routes Carbon Fluxes to Fuel Cell Cycle Progression. Mol Cell 62(4),532–545. doi: 10.1016/j.molcel.2016.02.017.

Galbraith, M.D., Andrysik, Z., Pandey, A., Hoh, M., Bonner, E.A., Hill, A.A., et al. (2017). CDK8 Kinase Activity Promotes Glycolysis. Cell Reports 21(6),1495–1506. doi: 10.1016/j.celrep.2017.10.058.

Gasch, A.P., and Werner-Washburne, M. (2002). The genomics of yeast responses to environmental stress and starvation. Funct Integr Genomics 2(4-5),181–192. doi: 10.1007/s10142-002-0058-2.

Ghillebert, R., Swinnen, E., Wen, J., Vandesteene, L., Ramon, M., Norga, K., et al. (2011). The AMPK/SNF1/SnRK1 fuel gauge and energy regulator: structure, function and regulation. Febs Journal 278(21),3978–3990. doi: 10.1111/j.1742-4658.2011.08315.x.

Godfrey, M., Touati, S.A., Kataria, M., Jones, A., Snijders, A.P., and Uhlmann, F. (2017). PP2A(Cdc55) Phosphatase Imposes Ordered Cell-Cycle Phosphorylation by Opposing Threonine Phosphorylation. Mol Cell 65(3),393–402 e393. doi: 10.1016/j.molcel.2016.12.018.

Goranov, A.I., and Amon, A. (2010). Growth and division--not a one-way road. Curr Opin Cell Biol 22(6),795–800. doi: 10.1016/j.ceb.2010.06.004.

Hardie, D.G. (2011). AMP-activated protein kinase-an energy sensor that regulates all aspects of cell function. Genes & Development 25(18),1895–1908. doi: 10.1101/gad.17420111.

Hartwell, L.H., Culotti, J., Pringle, J.R., and Reid, B.J. (1974). Genetic control of the cell division cycle in yeast. Science 183(4120),46–51.

Holt, L.J., Tuch, B.B., Villen, J., Johnson, A.D., Gygi, S.P., and Morgan, D.O. (2009). Global analysis of Cdk1 substrate phosphorylation sites provides insights into evolution. Science 325(5948),1682–1686. doi: 10.1126/science.1172867.

Icreverzi, A., de la Cruz, A.F., Van Voorhies, W.A., and Edgar, B.A. (2012). Drosophila cyclin D/Cdk4 regulates mitochondrial biogenesis and aging and sensitizes animals to hypoxic stress. Cell Cycle 11(3),554–568. doi: 10.4161/cc.11.3.19062.

Johnston, G.C., Pringle, J.R., and Hartwell, L.H. (1977). Coordination of Growth with Cell-Division in Yeast Saccharomyces-Cerevisiae. Experimental Cell Research 105(1),79–98. doi: Doi 10.1016/0014-4827(77)90154-9.

Kataria, M., Mouilleron, S., Seo, M.H., Corbi-Verge, C., Kim, P.M., and Uhlmann, F. (2018). A PxL motif promotes timely cell cycle substrate dephosphorylation by the Cdc14 phosphatase. Nat Struct Mol Biol 25(12),1093–1102. doi: 10.1038/s41594-018-0152-3.

Kurat, C.F., Wolinski, H., Petschnigg, J., Kaluarachchi, S., Andrews, B., Nafter, K., et al. (2009). Cdk1/Cdc28-Dependent Activation of the Major Triacylglycerol Lipase Tgl4 in Yeast Links Lipolysis to Cell-Cycle Progression. Molecular Cell 33(1),53–63. doi: 10.1016/j.molcel.2008.12.019.

Lowdon, M., and Vitols, E. (1973). Ribonucleotide reductase activity during the cell cycle of Saccharomyces cerevisiae. Arch Biochem Biophys 158(1),177–184. doi: 10.1016/0003-9861(73)90611-5.

Ma, X.Y., Wang, L., Huang, D., Li, Y.Y., Yang, D.D., Li, T.T., et al. (2017). Polo-like kinase 1 coordinates biosynthesis during cell cycle progression by directly activating pentose phosphate pathway. Nature Communications 8. doi: ARTN 150610.1038/s41467-017-01647-5.

McAlister, G.C., Nusinow, D.P., Jedrychowski, M.P., Wuhr, M., Huttlin, E.L., Erickson, B.K., et al. (2014). MultiNotch MS3 Enables Accurate, Sensitive, and Multiplexed Detection of Differential Expression across Cancer Cell Line Proteomes. Analytical Chemistry 86(14),7150–7158. doi: 10.1021/ac502040v.

Mochida, S., and Hunt, T. (2012). Protein phosphatases and their regulation in the control of mitosis. Embo Reports 13(3),197–203. doi: 10.1038/embor.2011.263.

Mok, J., Kim, P.M., Lam, H.Y., Piccirillo, S., Zhou, X., Jeschke, G.R., et al. (2010). Deciphering protein kinase specificity through large-scale analysis of yeast phosphorylation site motifs. Sci Signal 3(109),ra12. doi: 10.1126/scisignal.2000482.

Morgan, D.O. (ed.). (2007). The Cell Cycle: Principles of Control. London: New Science Press.

Morgan, D.O. (2008). SnapShot: cell-cycle regulators I. Cell 135(4),764–764 e761. doi: S0092-8674(08)01376-7 [pii] 10.1016/j.cell.2008.10.039.

Muller, D., Exler, S., Aguilera-Vazquez, L., Guerrero-Martin, E., and Reuss, M. (2003). Cyclic AMP mediates the cell cycle dynamics of energy metabolism in Saccharomyces cerevisiae. Yeast 20(4),351–367. doi: 10.1002/yea.967.

Nicastro, R., Tripodi, F., Gaggini, M., Castoldi, A., Reghellin, V., Nonnis, S., et al. (2015). Snf1 Phosphorylates Adenylate Cyclase and Negatively Regulates Protein Kinase A-dependent Transcription in Saccharomyces cerevisiae. Journal of Biological Chemistry 290(41),24715–24726. doi: 10.1074/jbc.M115.658005.

Oliveira, A.P., Ludwig, C., Picotti, P., Kogadeeva, M., Aebersold, R., and Sauer, U. (2012). Regulation of yeast central metabolism by enzyme phosphorylation. Molecular Systems Biology 8. doi: ARTN 62310.1038/msb.2012.55.

Ottoz, D.S., Rudolf, F., and Stelling, J. (2014). Inducible, tightly regulated and growth condition-independent transcription factor in Saccharomyces cerevisiae. Nucleic Acids Res 42(17),e130. doi: 10.1093/nar/gku616.

Pearce, E.J., and Pearce, E.L. (2018). Driving immunity: all roads lead to metabolism. Nature Reviews Immunology 18(2),81–+. doi: 10.1038/nri.2017.139.

Perez-Riverol, Y., Csordas, A., Bai, J.W., Bernal-Llinares, M., Hewapathirana, S., Kundu, D.J., et al. (2019). The PRIDE database and related tools and resources in 2019: improving support for quantification data. Nucleic Acids Research 47(D1),D442–D450. doi: 10.1093/nar/gky1106.

Pringle, J.R., and Hartwell, L.H. (1981). The Saccharomyces cerevisiae Cell Cycle.

Ptacek, J., Devgan, G., Michaud, G., Zhu, H., Zhu, X., Fasolo, J., et al. (2005). Global analysis of protein phosphorylation in yeast. Nature 438(7068),679–684. doi: 10.1038/nature04187.

Ramirez-Gaona, M., Marcu, A., Pon, A., Guo, A.C., Sajed, T., Wishart, N.A., et al. (2017). YMDB 2.0: a significantly expanded version of the yeast metabolome database. Nucleic Acids Research 45(D1),D440–D445. doi: 10.1093/nar/gkw1058.

Rogers, S., McCloy, R., Watkins, D.N., and Burgess, A. (2016). Mechanisms regulating phosphatase specificity and the removal of individual phosphorylation sites during mitotic exit. Bioessays 38,S24–S32. doi: 10.1002/bies.201670905.

Rosebrock, A.P. (2017). Methods for Synchronization and Analysis of the Budding Yeast Cell Cycle. Cold Spring Harb Protoc 2017(1),pdb top080630. doi: 10.1101/pdb.top080630.

Schwartz, D., and Gygi, S.P. (2005). An iterative statistical approach to the identification of protein phosphorylation motifs from large-scale data sets. Nat Biotechnol 23(11),1391–1398. doi: 10.1038/nbt1146.

Smith, F.C., Davies, S.P., Wilson, W.A., Carling, D., and Hardie, D.G. (1999). The SNF1 kinase complex from Saccharomyces cerevisiae phosphorylates the transcriptional repressor protein Mig1p in vitro at four sites within or near regulatory domain 1. Febs Letters 453(1-2),219–223. doi:Doi 10.1016/S0014-5793(99)00725-5.

Solaki, M., and Ewald, J.C. (2018). Fueling the Cycle: CDKs in Carbon and Energy Metabolism. Frontiers in Cell and Developmental Biology 6. doi: UNSP 9310.3389/fcell.2018.00093.

Swaffer, M.P., Jones, A.W., Flynn, H.R., Snijders, A.P., and Nurse, P. (2016). CDK Substrate Phosphorylation and Ordering the Cell Cycle. Cell 167(7),1750–1761 e1716. doi: 10.1016/j.cell.2016.11.034.

Tokiwa, G., Tyers, M., Volpe, T., and Futcher, B. (1994). Inhibition of G1 cyclin activity by the Ras/cAMP pathway in yeast. Nature 371(6495),342–345. doi: 10.1038/371342a0.

Touati, S.A., Kataria, M., Jones, A.W., Snijders, A.P., and Uhlmann, F. (2018). Phosphoproteome dynamics during mitotic exit in budding yeast. EMBO J 37(10). doi: 10.15252/embj.201798745.

Touati, S.A., and Uhlmann, F. (2018). A global view of substrate phosphorylation and dephosphorylation during budding yeast mitotic exit. Microb Cell 5(8),389–392. doi: 10.15698/mic2018.08.644.

Tu, J.L., and Carlson, M. (1995). Reg1 Binds to Protein Phosphatase Type-1 and Regulates Glucose Repression in Saccharomyces-Cerevisiae. Embo Journal 14(23),5939–5946. doi: DOI 10.1002/j.1460-2075.1995.tb00282.x.

Tudzarova, S., Colombo, S.L., Stoeber, K., Carcamo, S., Williams, G.H., and Moncada, S. (2011). Two ubiquitin ligases, APC/C-Cdh1 and SKP1-CUL1-F (SCF)-beta-TrCP, sequentially regulate glycolysis during the cell cycle. Proceedings of the National Academy of Sciences of the United States of America 108(13),5278–5283. doi: 10.1073/pnas.1102247108.

Tyanova, S., Temu, T., Sinitcyn, P., Carlson, A., Hein, M.Y., Geiger, T., et al. (2016). The Perseus computational platform for comprehensive analysis of (prote)omics data. Nature Methods 13(9),731–740. doi: 10.1038/Nmeth.3901.

Ubersax, J.A., Woodbury, E.L., Quang, P.N., Paraz, M., Blethrow, J.D., Shah, K., et al. (2003). Targets of the cyclin-dependent kinase Cdk1. Nature 425(6960),859–864. doi: 10.1038/nature02062.

Vander Heiden, M.G., and DeBerardinis, R.J. (2017). Understanding the Intersections between Metabolism and Cancer Biology. Cell 168(4). doi: 10.1016/j.cell.2016.12.039.

Vaupel, P., Schmidberger, H., and Mayer, A. (2019). The Warburg effect: essential part of metabolic reprogramming and central contributor to cancer progression. International Journal of Radiation Biology 95(7),912–919. doi: 10.1080/09553002.2019.1589653.

Verbinnen, I., Ferreira, M., and Bollen, M. (2017). Biogenesis and activity regulation of protein phosphatase 1 (vol 45, pg 89, 2017). Biochemical Society Transactions 45,583–584. doi: 10.1042/Bst-2016-0154c_Cor.

Wang, H.Z., Nicolay, B.N., Chick, J.M., Gao, X.L., Geng, Y., Ren, H., et al. (2017). The metabolic function of cyclin D3-CDK6 kinase in cancer cell survival. Nature 546(7658),426–+. doi: 10.1038/nature22797.

Yang, F., Shen, Y., Camp, D.G., 2nd, and Smith, R.D. (2012). High-pH reversed-phase chromatography with fraction concatenation for 2D proteomic analysis. Expert Rev Proteomics 9(2),129–134. doi: 10.1586/epr.12.15.

Zhang, J., Zhao, J., Dahan, P., Lu, V., Zhang, C., Li, H., et al. (2018). Metabolism in Pluripotent Stem Cells and Early Mammalian Development. Cell Metabolism 27(2),332–338. doi: 10.1016/j.cmet.2018.01.008.

Zhang, L.C., and Elias, J.E. (2017). Relative Protein Quantification Using Tandem Mass Tag Mass Spectrometry. Proteomics: Methods and Protocols 1550,185–198. doi: 10.1007/978-1-4939-6747-6_14.

Zhao, G., Chen, Y., Carey, L., and Futcher, B. (2016). Cyclin-Dependent Kinase Co-Ordinates Carbohydrate Metabolism and Cell Cycle in S. cerevisiae. Mol Cell 62(4),546–557. doi: 10.1016/j.molcel.2016.04.026.

