## Supplementary Figures for "Multiple layers of phospho-regulation coordinate metabolism and the cell cycle in budding yeast"

**Supplementary Tables**

Supplementary Table 1: Excel file containing total proteome data

Supplementary Table 2: Excel file containing processed and raw data for phosphoproteome

Supplementary Table 3: Excel file containing overview of all determined phosphosite-metabolite correlations (see Figure 4 and Supplementary Figure 3)

**Supplementary Figures**

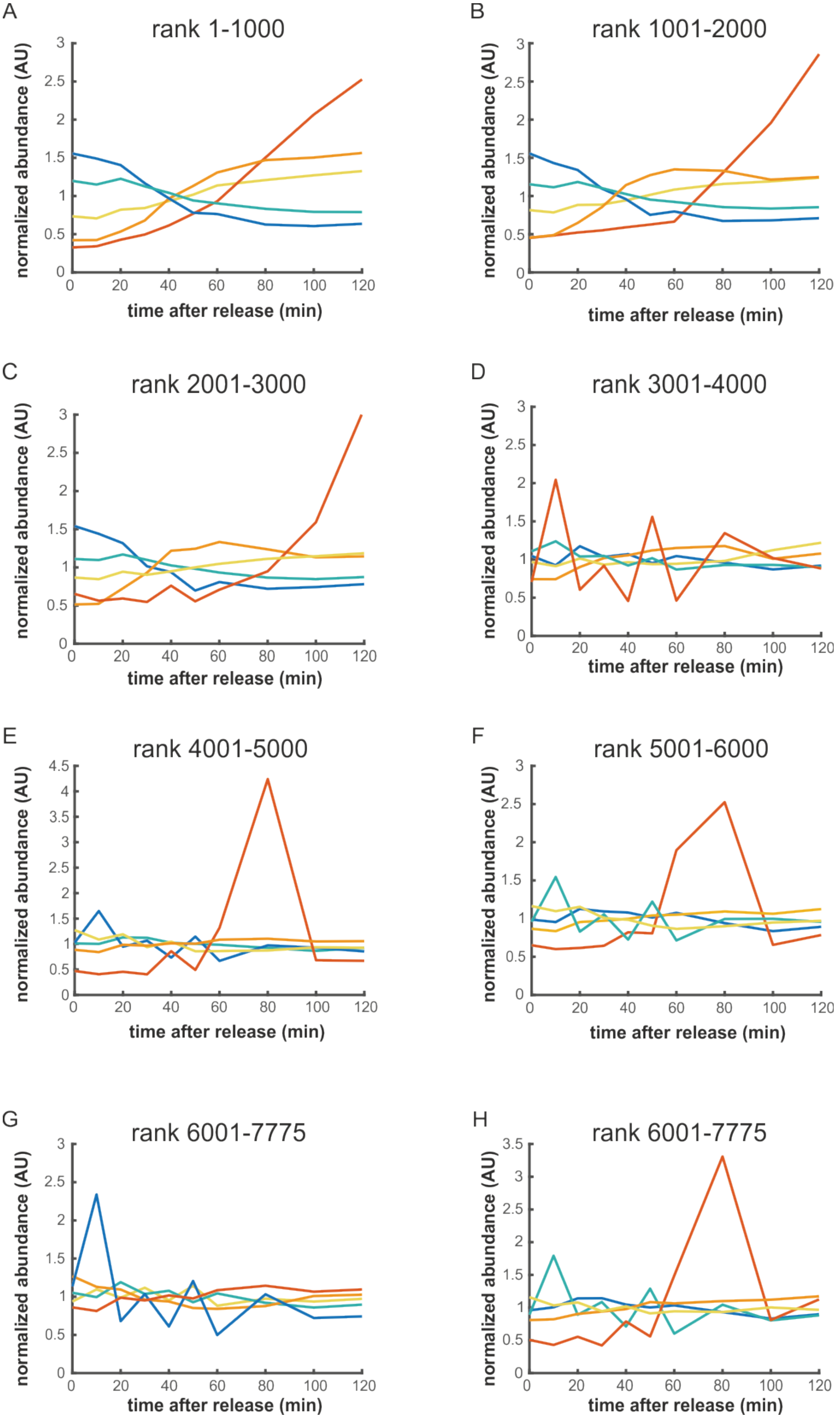
A
rank 1-1000
B
rank 1001-2000
C
rank 2001-3000
D
rank 3001-4000
E
rank 4001-5000
F
rank 5001-6000
G
rank 6001-7775
H
rank 6001-7775
normalized abundance (AU)
time after release (min)
0
20
40
60
80
100
120
0
0.5
1
1.5
2
2.5
3
3.5
4
4.5

I

J

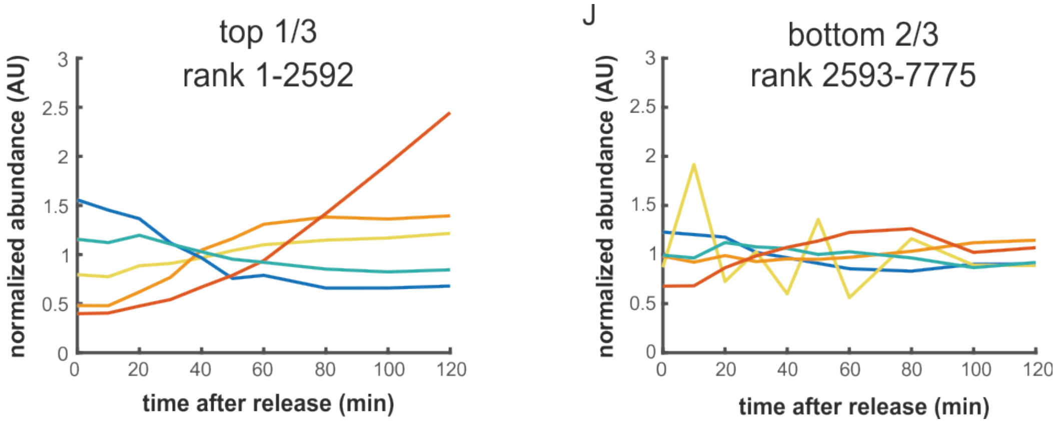


**Supplementary Figure 1:** To empirically confirm that our statistical ranking of phosphosites (see methods) separates changing from non-changing time courses, we performed K-means clustering (see methods) on sets of 1000 phosphosites of ascending rank (A-H). Plotted are the means of each of the five clusters. We note that the red clusters with large peaks at t=80 minutes in E, F, and H only correspond to three to four phosphosites each. There seems to be technical noise in 10-20 phosphopeptides only in replicate 1 at time point 80 minutes.  Based on these comparisons we set the cut-off for further analysis at the top third, rank 2592. Panels I and J show the comparison of the averages of the k-means clustered time profiles for the top third and bottom two thirds ranking sites.

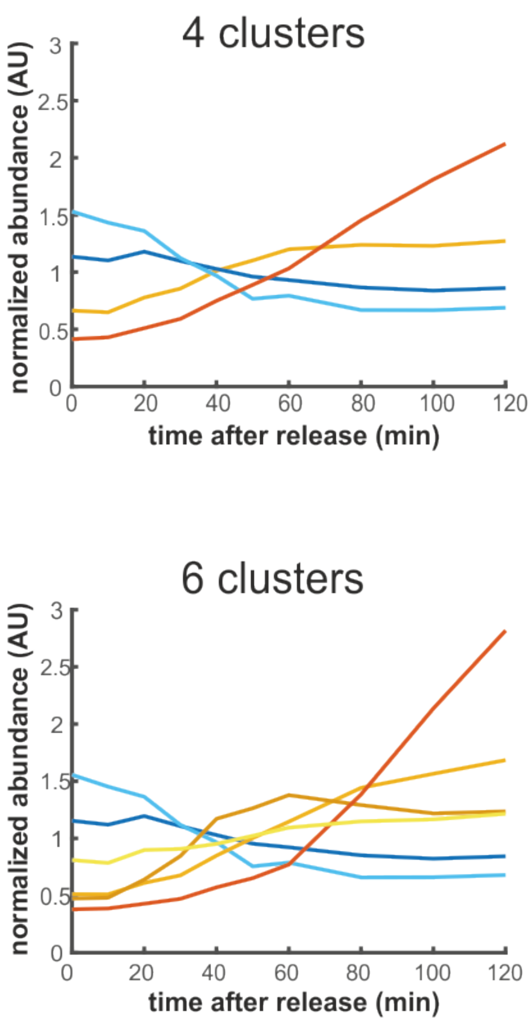


**Supplementary Figure 2**: Results of k-means clustering of the top third ranking sites with the number of clusters set to four, five, six, or eight. Plotted are the cluster averages. Since all four settings give qualitatively similar results, we chose five clusters as a balance between cluster size and resolution.

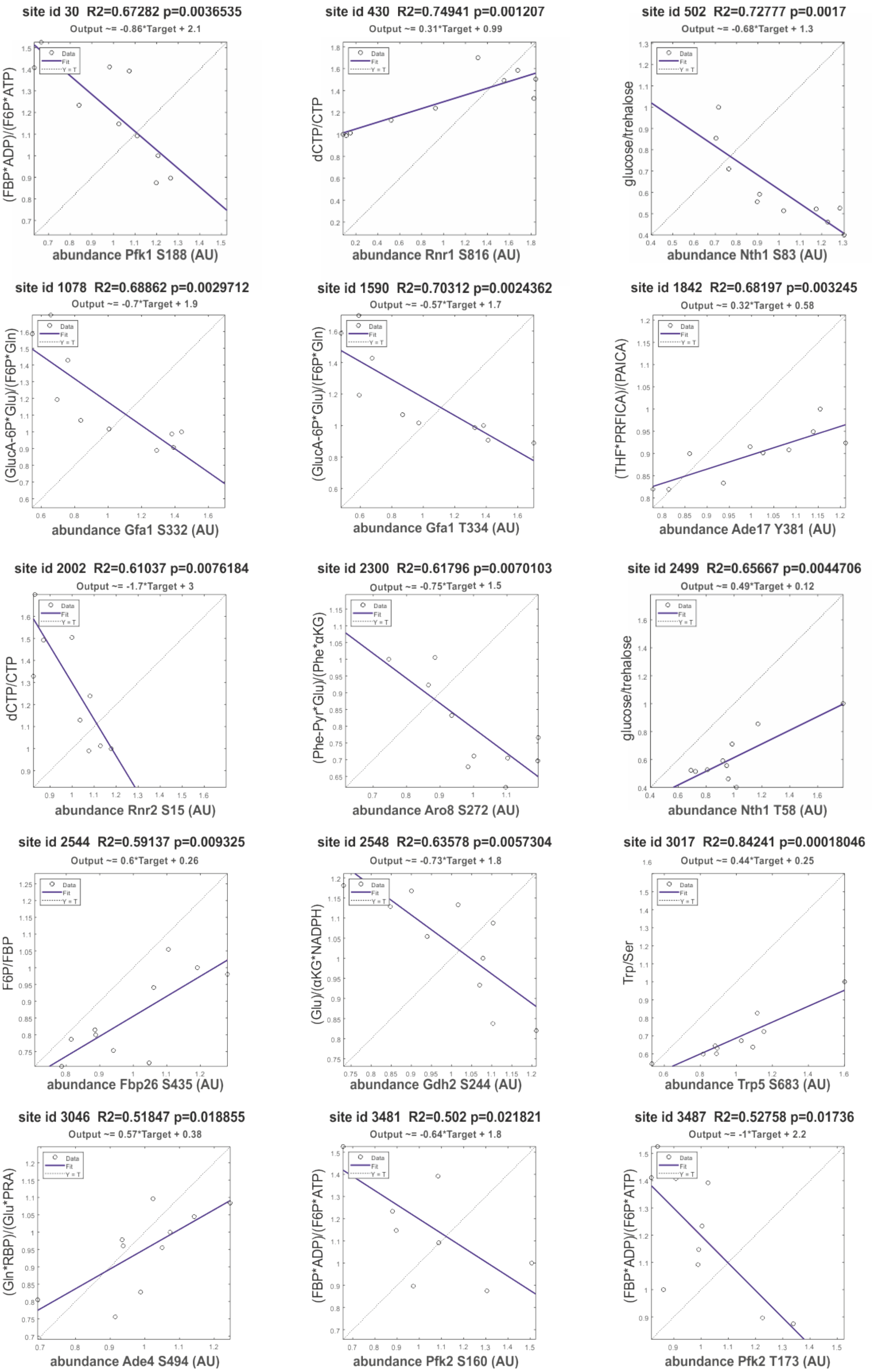
site id 30 R2=0.67282 p=0.0036535
Output ~= -0.86*Target + 2.1
(FBP*ADP)/(F6P*ATP)
abundance Pfk1 S188 (AU)
site id 430 R2=0.74941 p=0.001207
Output ~= 0.31*Target + 0.99
dCTP/CTP
abundance Rnr1 S816 (AU)
site id 502 R2=0.72777 p=0.0017
Output ~= -0.68*Target + 1.3
glucose/trehalose
abundance Nth1 S83 (AU)
site id 1078 R2=0.68862 p=0.0029712
Output ~= -0.7*Target + 1.9
(GlucA-6P*Glu)/(F6P*Gln)
abundance Gfa1 S332 (AU)
site id 1590 R2=0.70312 p=0.0024362
Output ~= -0.57*Target + 1.7
(GlucA-6P*Glu)/(F6P*Gln)
abundance Gfa1 T334 (AU)
site id 1842 R2=0.68197 p=0.003245
Output ~= 0.32*Target + 0.58
(THF*PRFICA)/(PAICA)
abundance Ade17 Y381 (AU)
site id 2002 R2=0.61037 p=0.0076184
Output ~= -1.7*Target + 3
dCTP/CTP
abundance Rnr2 S15 (AU)
site id 2300 R2=0.61796 p=0.0070103
Output ~= -0.75*Target + 1.5
(Phe-Pyr*Glu)/(Phe*αKG)
abundance Aro8 S272 (AU)
site id 2499 R2=0.65667 p=0.0044706
Output ~= 0.49*Target + 0.12
glucose/trehalose
abundance Nth1 T58 (AU)
site id 2544 R2=0.59137 p=0.009325
Output ~= 0.6*Target + 0.26
F6P/FBP
abundance Fbp26 S435 (AU)
site id 2548 R2=0.63578 p=0.0057304
Output ~= -0.73*Target + 1.8
(Glu)/(αKG*NADPH)
abundance Gdh2 S244 (AU)
site id 3017 R2=0.84241 p=0.00018046
Output ~= 0.44*Target + 0.25
Trp/Ser
abundance Trp5 S683 (AU)
site id 3046 R2=0.51847 p=0.018855
Output ~= 0.57*Target + 0.38
(Gln*RBP)/(Glu*PRA)
abundance Ade4 S494 (AU)
site id 3481 R2=0.502 p=0.021821
Output ~= -0.64*Target + 1.8
(FBP*ADP)/(F6P*ATP)
abundance Pfk2 S160 (AU)
site id 3487 R2=0.52758 p=0.01736
Output ~= -1*Target + 2.2
(FBP*ADP)/(F6P*ATP)
abundance Pfk2 T173 (AU)
Data
Fit
Y = T

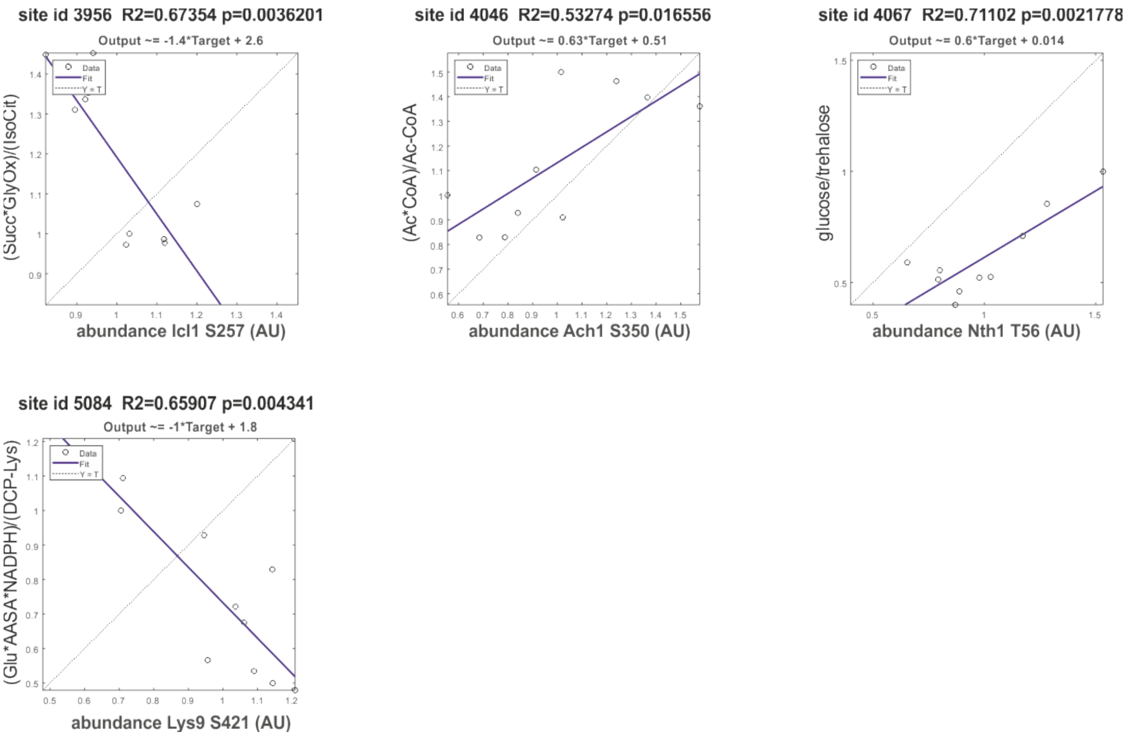


**Supplementary Figure 3:** Phosphorylation sites on metabolic enzymes with putative regulatory function. We correlated the normalized abundances of phosphosites on metabolic enzymes with the corresponding product to substrate ratios (data from (Ewald et al., 2016)). We show all correlations with an $R^2$ greater than 0.5, the results of all available data are reported in Supplementary Table 3. Uncommon metabolite abbreviations: GlucA-6P: D-Glucosamine 6-phosphate ; PRFICA: 1-(5'-Phosphoribosyl)-5-formamido-4-imidazolecarboxamide; PAICA: 1-(5'-Phosphoribosyl)-5-amino-4-imidazolecarboxamide; RBP: 5-Phospho-alpha-D-ribose 1-diphosphate;PRA: 5-Phosphoribosylamine ; AASA: L-2-Aminoadipate 6-semialdehyde ; DCP-Lys: N6-(L-1,3-Dicarboxypropyl)-L-lysine ; (see Supplementary Table 3 for further information on metabolites and the catalysed reactions)

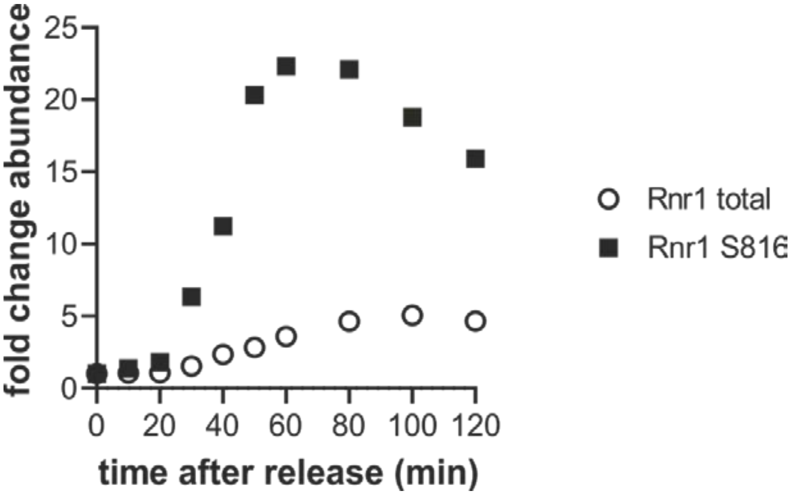


**Supplementary Figure 4:** Rnr1 phosphorylation increases more strongly than total Rnr1 protein abundance. Comparison between the fold increase in S816 (black squares) as determined from the phospho-enriched samples versus the Rnr1 protein (white circles) as determined from the total proteome samples.

A B

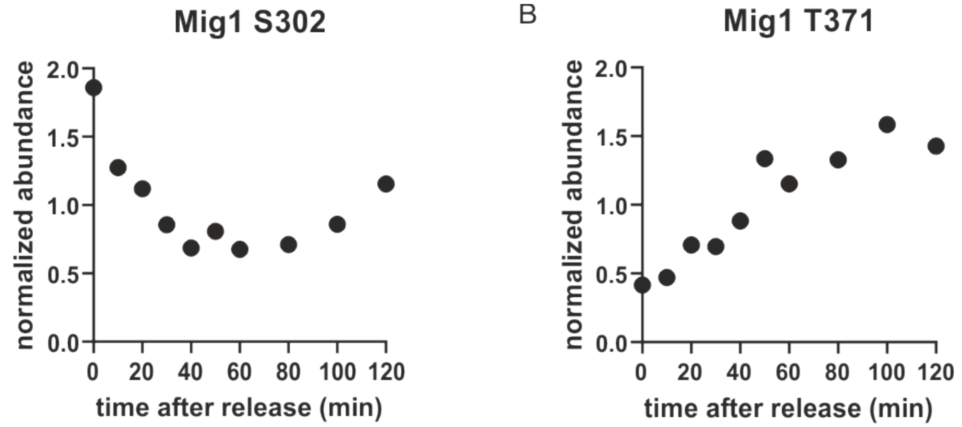


C Mig1 regulatory domain

... 210
ALSSLSNSHS GSRLKLNALS SLQMMTPIAS SAPRTVFIDG PEQKQLQQQQ NSLSPRYSNT
VILPRPRSLT DFQGLNNANP NNNGSLRAQT QSSVQLKRPS SVLSLNDLLV GQRNTNESDS
DFTTGGEDEE DGLKDPSNSS IDNLEQDYLQ EQSRKKSKTS TPTTMLSRST SGTNLHTLGY
VMNQNHLHFS SSSPDFQKEL 410 ...

**Supplementary Figure 5:** A. Mig1 S302 is dephosphorylated in a similar pattern as Snf1. (This site was only detected in replicate 1) B. Mig1 T371 is a proline-directed site that increases during the cell cycle (This site was only detected in replicate 2) C. Amino acid sequence of the Mig1 regulatory domain containing known Snf1 sites and cell cycle regulated sites. Red: Known Snf1 phosphorylation sites. Blue: Sites whose phosphorylation changes through the cell cycle in this study (see A and B). Green: Previously identified CDK target site (Holt et al., 2009); Grey: Other phosphorylation sites annotated on BioGrid (Oughtred et al., 2019) with unknown functions.

### Supplementary references:

Ewald, J.C., Kuehne, A., Zamboni, N., and Skotheim, J.M. (2016). The Yeast Cyclin-Dependent Kinase Routes Carbon Fluxes to Fuel Cell Cycle Progression. *Mol Cell* 62(4), 532-545. doi: 10.1016/j.molcel.2016.02.017.

Holt, L.J., Tuch, B.B., Villen, J., Johnson, A.D., Gygi, S.P., and Morgan, D.O. (2009). Global analysis of Cdk1 substrate phosphorylation sites provides insights into evolution. *Science* 325(5948), 1682-1686. doi: 10.1126/science.1172867.

Oughtred, R., Stark, C., Breitkreutz, B.J., Rust, J., Boucher, L., Chang, C., et al. (2019). The BioGRID interaction database: 2019 update. *Nucleic Acids Research* 47(D1), D529-D541. doi: 10.1093/nar/gky1079.
